# Shared and context-dependent cortico–striatal–cerebellar organization of temporal prediction

**DOI:** 10.64898/2026.11.21.768330

**Authors:** Ana Luísa Pinho, Velda Koranteng-Addo, Anmar Alsibaie, Nidhi Patel, Jörn Diedrichsen, Jessica A. Grahn

## Abstract

Temporal prediction is thought to draw on two subcortical structures: the basal ganglia for the periodic beat, and the cerebellum for the aperiodic interval. Lesion, imaging and electrophysiological studies converge on this division, yet each tests it within a single modality and task, leaving its generality unknown. We developed a battery of timing tasks for multi-task fMRI, densely sampling 31 non-clinical adults in every combination of temporal structure, sensory modality and task. Encoding engaged the same cortico–striatal–cerebellar architecture throughout. Within it, the dorsal striatum was uniquely selective for the production task. Pooled across tasks, it responded more to auditory beat than interval sequences, whereas the cerebellum did not distinguish them. Yet, removing regularity did not lower that response: the preference is for the beat, not for predictability. Temporal prediction therefore draws on a shared architecture whose striatal node is context-dependent: tuned to periodic time, and to time turned into action.

## Main

Temporal prediction is the anticipation of when future events will occur [1]. It underpins attention, perception and motor coordination [2], and draws on regularities extracted from past experience: periodic streams that create a rhythmic structure, or aperiodic streams in which a specific duration must be learned through exposure [3]. Two subcortical structures are central to this ability [4]: the basal ganglia support periodic, beatbased prediction, and the cerebellum supports aperiodic, interval-based prediction [3, 5, 6]. Whether their contributions are coordinated [7] or computationally and neurally separable [3, 5, 6] remains debated.

The strongest support for separation is causal, and comes from a double dissociation: patients with cerebellar degeneration lose the benefit of interval-based prediction but retain beat-based prediction, whereas those with Parkinson’s disease show the reverse pattern [3]. Consistent with this dissociation, Parkinson’s patients are also selectively impaired at discriminating beat-based rhythms [8]. Lesions to either structure weaken neural entrainment to a regular auditory rhythm, but only cerebellar lesions reduce entrainment stability [9]. Functional imaging in healthy individuals supports this division into distinct circuits: beat-based timing recruits a striato-thalamo-cortical circuit, whereas interval-based timing recruits an olivocerebellar one [6]. Within the striatum, activity reflects prediction of the beat rather than its detection [10].

However, attributing the observed neural dissociations to beat-based versus interval-based structure is difficult, because the supporting studies differ in several other dimensions. First, they have typically used a single sensory modality, leaving the proposed division untested across audition and vision within a single design. Second, the required response has varied from producing a duration to judging one. Because this distinction alone alters the timing circuits engaged [11], differences ascribed to beat versus interval timing may instead reflect differences in response demands. Finally, the evidence spans two forms of timing with distinct neural substrates [12]: explicit timing, in which duration is directly judged, and implicit timing, in which duration is never reported but regular temporal structure may be used to predict upcoming events and optimize prospective behaviour.

The basal ganglia and cerebellum are also embedded in integrated but partially separable thalamo-cortical loops with the premotor and prefrontal cortex [4, 13, 14]. Their engagement is therefore not strictly determined by anatomy. For example, the cerebellum shapes anticipatory cortical dynamics differently depending on the predictive context [15]. How cortical engagement of these circuits varies across sensory modality, task and temporal structure remains largely uncharacterized [9].

To disentangle these factors, we developed a battery of duration-based timing tasks crossing three dimensions in the same participants: the temporal structure of the events (beat *vs.* interval), the sensory modality (auditory *vs.* visual), and the timing behaviour required by each task. Every trial began with an *encoding* period, in which participants encoded the timing of a sequence of events marking two repetitions of a *standard* duration, followed by a response period (Figure 1). Previous studies used isochronous sequences [3, 6], a special case of rhythm in which a single duration repeats throughout. Here, the two temporal structures differed only in how the standards were spaced: at integer ratios implying a periodic pulse in the beat condition [16, 17], and irregularly in the interval condition. With the standard presented equally often in both conditions, periodicity was dissociated from the repetition of a single duration. The response period defined the task: production and perception both call for an explicit temporal judgement, one by reproducing a duration and the other by discriminating it. *Non-Temporal Feature Discrimination* (NTFD) instead calls for a decision about a non-temporal feature, engaging timing only implicitly to anticipate the judged event and optimize performance [12, 18]. Because encoding was matched across tasks while the response varied, this design isolates the formation of temporal predictions.

**Figure 1.**
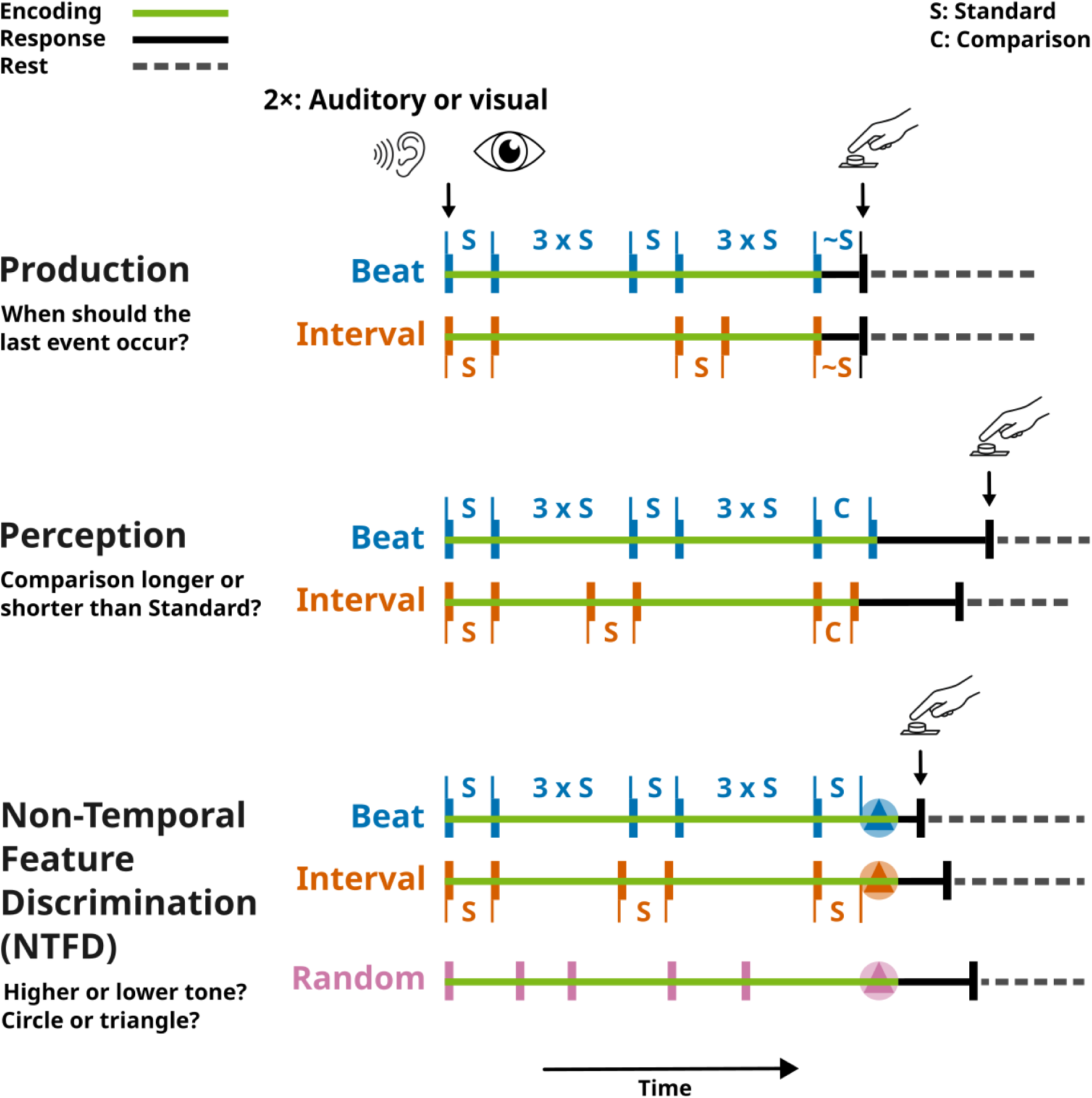
Schematic representation of trials for each task: production, perception, and Non-Temporal Feature Discrimination. Each paradigm was tested with blocks of trials in the auditory and the visual modality. Standards (S) on each trial were selected from one of five durations: 459 ms, 510 ms, 561 ms, 612 ms, and 663 ms. During the encoding period (solid green line), participants attended to a sequence of 80 ms-long events (rectangular bars) presented either periodically (beat, in blue) or aperiodically (interval, in orange). In the *beat* condition, consecutive standards were separated by three times the standard (3×S), so that all event onsets fell on a regular grid, whereas in the *interval* condition, they were separated by irregular durations, preventing the establishment of a beat percept. Events were 440 Hz beeps in the auditory modality or rectangle-shaped flashes in the visual modality. The response period (solid black line) began at the end of encoding and ended with the button press (black tick), made according to the requirement of each task. The remainder of the 2000 ms response window and the 2000 ms inter-trial interval formed the rest period (grey dashed line). For the production task, participants pressed the button when they felt the final event should occur, to produce a duration matching the standard. For the perception task, participants indicated whether the comparison (C) was longer or shorter than the standard. For the NTFD task, participants discriminated whether the final auditory beep was higher (880 Hz) or lower (220 Hz) than the preceding beeps, or whether the visual shape was a circle or a triangle. A variant of the NTFD task also included a *random* temporal structure, in which the timing of all encoding events was jittered.

To map the brain systems that form these predictions, we adapted a deep behavioural phenotyping frame-work to multi-task fMRI [19]. Sampling each participant densely across conditions separates their shared temporal-encoding response from condition-specific effects [20] and finely characterizes the regions involved [21]. It also supports individualized *Regions-of-Interest* (ROIs), which map function with greater sensitivity and precision than group-defined ROIs [22, 23].

Thirty-one adults with no clinical diagnosis completed all three tasks, in both modalities, over two imaging sessions; a variant of the NTFD task added unpredictable (random) sequences, with no temporal regularity. We hypothesized that the dorsal striatum would respond more to periodic (beat) sequences, and the cerebellum to aperiodic (interval) sequences, in which duration must be encoded in absolute terms. We find that temporal encoding engages a distributed cortico–striatal–cerebellar architecture, shared across tasks and sensory modalities. Within it, the dorsal striatum stands apart twice: it is selective for the production task, and it responds more to auditory beat than interval sequences, a difference the cerebellum does not show. Removing regularity does not lower this striatal response, so its preference is for the beat rather than for predictability. Temporal prediction therefore draws on a shared architecture whose striatal node depends on the task and on periodicity.

## Results

### Temporal Encoding Engages a Distributed Cortico–Striatal–Cerebellar Architecture

To identify regions supporting temporal encoding during predictive timing, we examined the encoding *vs.* rest contrast collapsed across all tasks, sensory modalities, and temporal structures. This yielded a group-level spatial map of where encoding-related activation was reliably greater than baseline, spanning cortico–striatal–cerebellar systems bilaterally. On the cortical surface (Figure 2a), it extended across the lateral and medial premotor cortices, comprising the *Pre-Supplementary Motor Area* (PreSMA), *Supplementary Motor Area* (SMA), *dorsal Premotor Area* (PMD) and *ventral Premotor Area* (PMV) [24], together with the posterior-dorsal section of the *Frontal Operculum* (FO). The sensory cortices showed modality-specific responses [25, 26]: in the auditory domain, activation was localized in *Heschl’s gyrus* and extended over the superior temporal plane and adjacent perisylvian cortices; in the visual domain, it was localized in both foveal and peripheral occipital cortex and in higher-order visual regions, including the *Superior Parietal Lobe* (SPL). Modality-specific maps are shown in Extended Data Fig. E1, and peak-level statistics in Extended Data Table E1.

**Figure 2.**
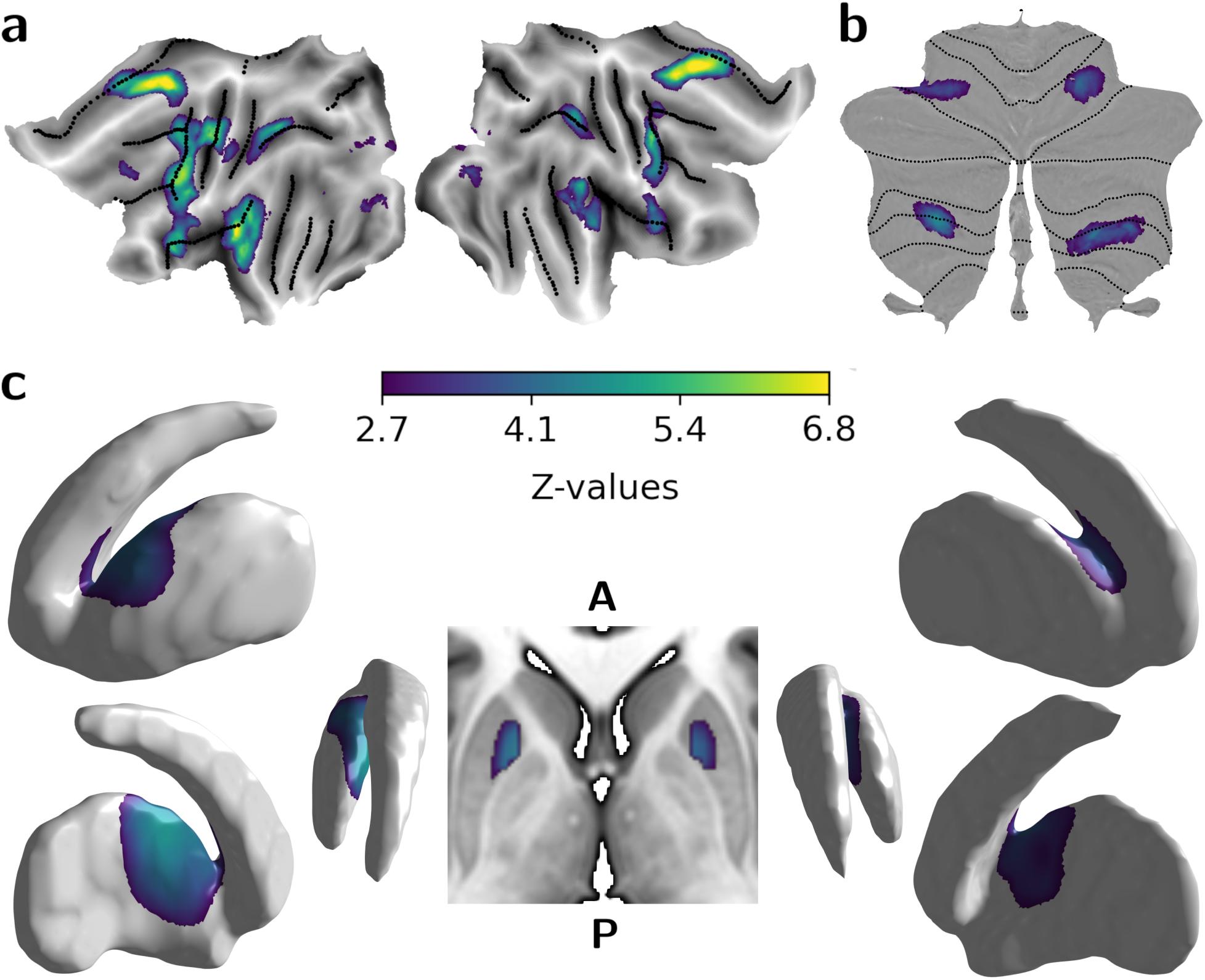
Group-level *Z*-maps for encoding *vs.* rest, collapsed across tasks, sensory modalities, and temporal structures, in cortical and subcortical regions. The display threshold (whole-brain FDR, *q ≤* 0.05) and colour bar limits were computed from the volume-based group map and applied to all panels. The contrast is shown in three coordinate spaces: (a) flat projection of activations computed on the fs LR32k surface, with left and right maps corresponding to the left and right cortical hemispheres; (b) flat projection of volume-based cerebellar activations from both hemispheres onto SUIT space; and (c) projection of volume-based striatal activations onto left and right rostrodorsal striatum surface meshes (left and right sections), with the centre section showing bilateral putaminal activation on an axial MNI152 slice at *z* = −0.23 mm.

In the cerebellum, encoding-related activations were observed bilaterally in lobules VI, VIIb, VIIIa, and Crus I [27]. According to a recently developed functional cerebellar atlas [28], these activations were predominantly localized in the D3 regions of the *Demand* domain and extended to the mouth (M2) and hand (M3) areas of the *Motor* domain (Figure 2b and Extended Data Table E1).

Subcortical contributions to temporal encoding were also evident in the dorsal striatum. Activations were located primarily in the anterior–medial section of the Putamen and extended to the head region of the Caudate Nucleus [29]. Responses were bilateral, with greater prominence in the left hemisphere (Figure 2c). Functional subdivisions reported in Extended Data Table E1 were defined by their structural connectivity with prefrontal association and premotor cortices [30].

Together, encoding-related activity spanned cortical, striatal and cerebellar regions, in a pattern that confirms previous findings [4, 6, 8, 11, 31].

### Temporal Encoding Combines Shared Representations across Behavioural Contexts with Task-Dependent Striatal Modulation

Having established that a cortico–striatal–cerebellar system was engaged during temporal encoding, we asked whether all these regions contributed in the same way, or whether their activation differed with sensory input, task demands, and periodicity. To account for inter-individual variability and improve sensitivity [22, 23], every region of the encoding map was delineated with an individualized ROI framework, with boundaries defined separately for each participant (spatial coverage in Extended Data Fig. E2; technical details in Methods). The auditory and visual cortices, which respond selectively to their own modality, provided a benchmark against which to assess the modality invariance of the other regions.

We quantified the *Percent Signal Change* (PSC) during the encoding period relative to the resting baseline in each of the 12 task conditions (2 sensory modalities, 3 tasks, 2 temporal structures). Inter-regional functional similarity was characterized by correlating these activation profiles (Figure 3a) and visualized with classical *Multidimensional Scaling* (MDS; see Methods), in which ROIs with similar functional profiles are positioned close together and dissimilar ones farther apart (Figure 3b). The first two dimensions separated a core set of premotor, cerebellar, and striatal regions from the sensory cortices (Extended Data Fig. E3a). A two-way *Repeated Measures Analysis-of-Variance* (RM ANOVA), with modality (auditory, visual) and task (production, perception, NTFD) as within-subject factors, revealed the expected modality-specific effects in the sensory cortices: auditory cortex showed greater activity for auditory tasks (Modality × Task: *F*_(2,60)_ = 14.26, *p* = 8.56 × 10^−6^; *ɛ* = 0.98), whereas the visual cortex showed greater activity for visual tasks (Modality × Task: *F*_(2,60)_ = 6.28, *p* = 0.003, *ɛ* = 0.98). In contrast, no other region showed a Modality × Task interaction (all: *F*_(2,60)_ ≤ 0.68, *p≥* 0.512), and only PMD showed a main effect of modality, with slightly greater activity for visual than auditory tasks (*F*_(1,30)_ = 9.93, *p* = 0.004, *ɛ* = 1; Figure 4).

**Figure 3.**
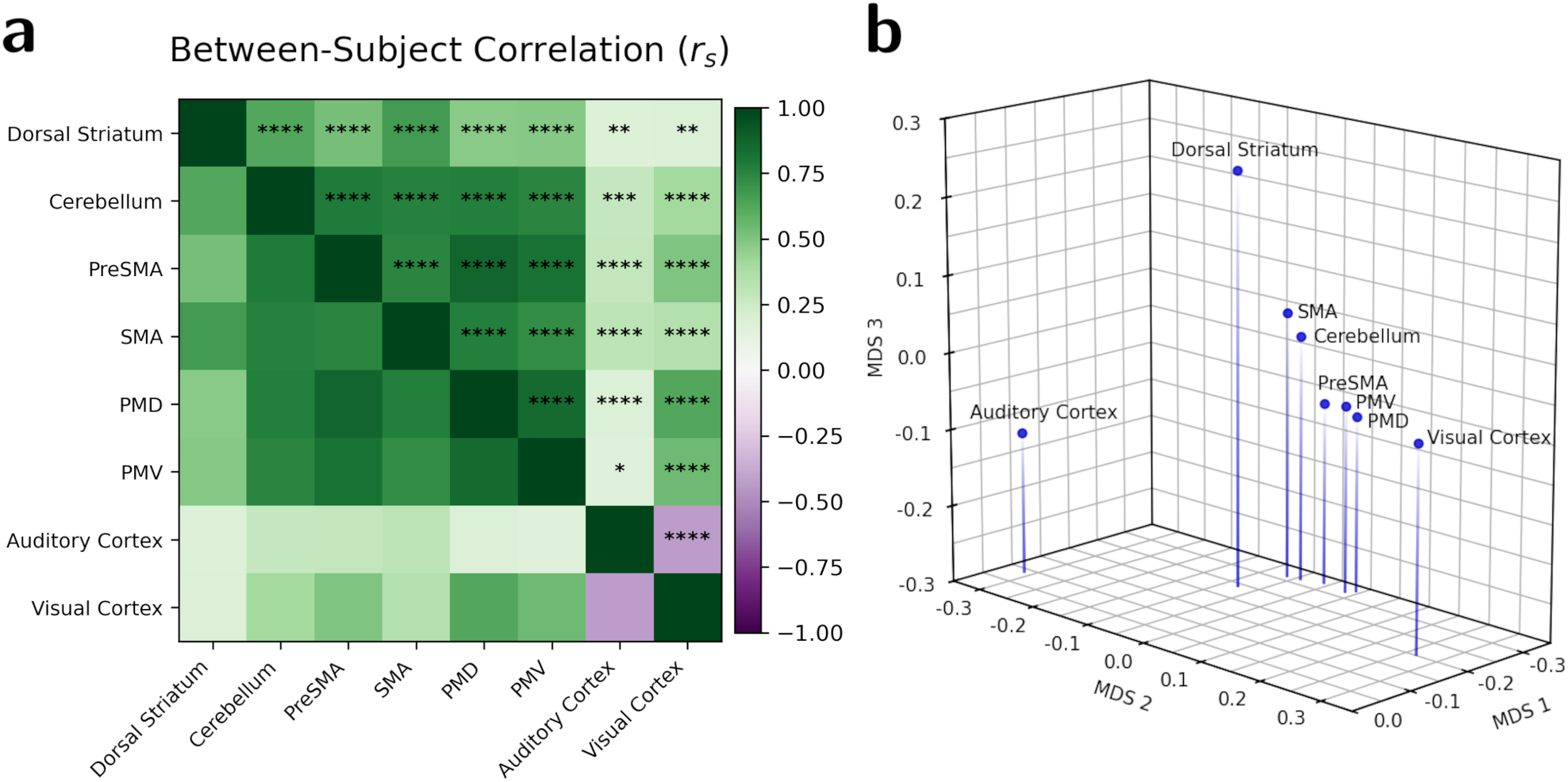
ROI-similarity structure during temporal encoding. **(a)** ROI-by-ROI similarity matrix computed from subjectwise correlations of condition-evoked PSC activation profiles. Each cell reports the mean *Pearson* correlation across subjects (*rs*), quantifying the extent to which activation profiles across experimental conditions covary between pairs of individualized ROIs. Stars denote the uncorrected *p*-value at which the Fisher-*Z*-transformed correlations differ significantly from zero across subjects; every ROI pair survived Holm–Bonferroni correction across the 28 pairs (*α*_FWE_ = 0.05; * *p <* 0.05, ** *p <* 0.01, *** *p <* 0.001, **** *p <* 10^−4^). **(b)** Three-dimensional classical MDS representation of the ROI similarity structure derived from the similarity matrix. Each point represents an ROI, and greater distances indicate more dissimilar activation profiles. Axes are ordered by decreasing eigenvalue magnitude.

**Figure 4.**
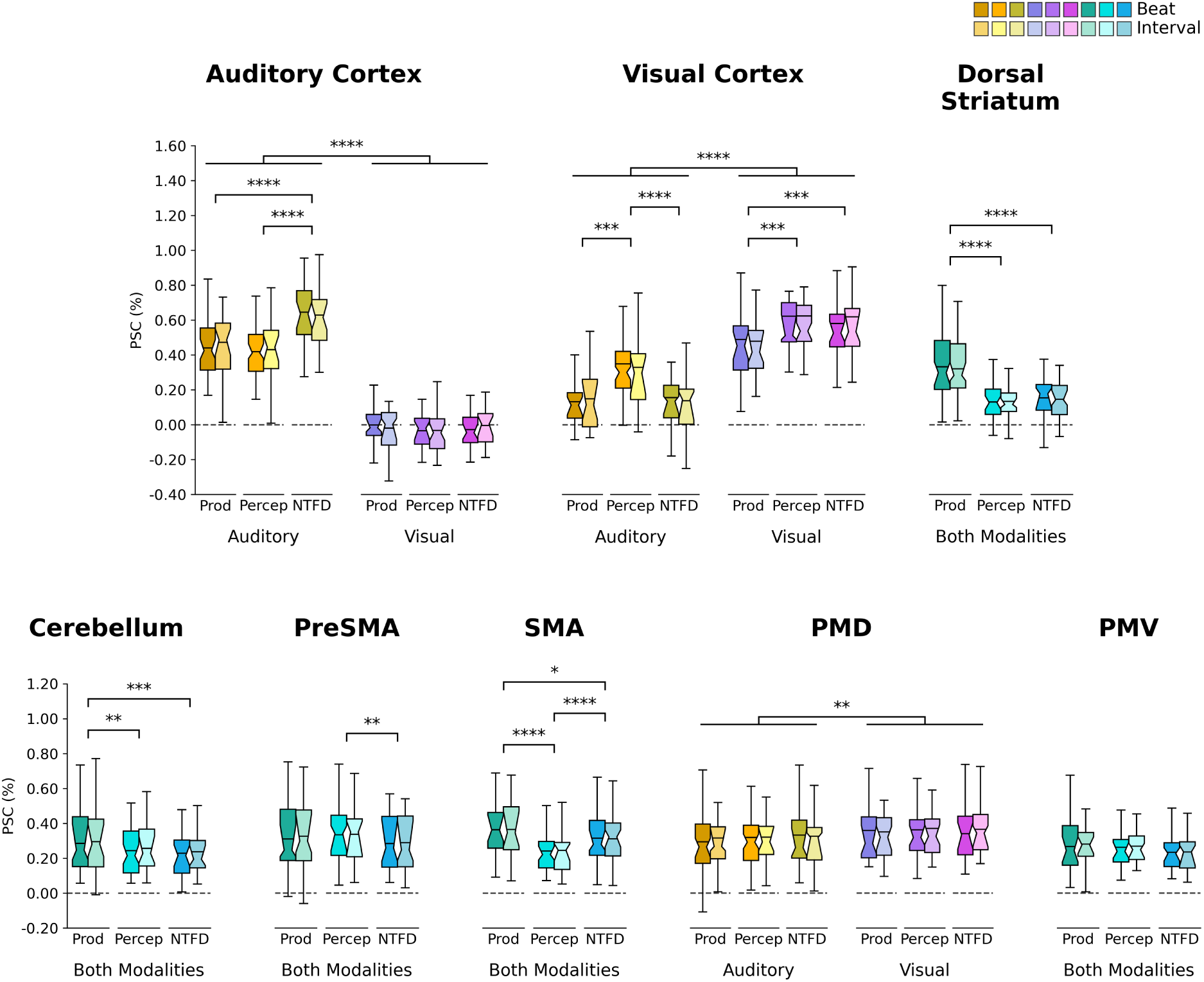
ROI-activation profiles during temporal encoding. Activation profiles are expressed as PSC relative to rest across sensory modalities, tasks, and temporal structures (beat and interval shown separately for visualization; analyses use PSC averaged across them). Boxplots show distributions for production (Prod), perception (Percep), and NTFD tasks; effects are assessed with two-way RM ANOVAs, with annotations indicating post hoc comparisons surviving omnibus and FWE-corrected testing (*α*_FWE_ = 0.05; * *p <* 0.05, ** *p <* 0.01, *** *p <* 0.001, **** *p <* 10^−4^). Plots averaged across modalities show task effects only; modality-specific plots reflect significant modality (lines spanning task groups) and, except in PMD, Modality × Task effects (pairwise annotations). Boxes represent the *Interquartile Range* (IQR), notches indicate the bootstrap 95% confidence interval of the median, whiskers extend to 1.5×IQR, and horizontal lines denote the mean. The dashed horizontal lines mark PSC = 0.

The third MDS dimension separated the dorsal striatum from the multimodal cortical regions and the cerebellum. This axis captured differences in task-specific modulation (Extended Data Fig. E3b and c). The dorsal striatum showed a strong effect of task (*F*_(2,60)_ = 45.13, *p* = 1.10 × 10^−12^, *ɛ* = 0.79), with the highest activity observed in the production task (Figure 4). These task differences did not depend on sensory modality (Modality × Task: *F*_(2,60)_ = 0.19, *p* = 0.826, *ɛ* = 0.94). Post hoc comparisons (Holm–Bonferroni: three taskpair tests, *α*_FWE_ = 0.05, smallest *p ≤* 0.05*/*3) revealed significantly higher encoding-related activation for production than for perception (*t*_(30)_ = 7.54, *p* = 2.07 × 10^−8^, *d* = 1.26 [Cohen’s d]) and for production than for NTFD (*t*_(30)_ = 7.07, *p* = 7.38 × 10^−8^, *d* = 1.10), representing the largest task-related effect sizes observed across all non-sensory regions examined.

In contrast to the dorsal striatum, no cortical region exhibited an equally pronounced selectivity for the production task. Encoding-related activity in PMD and PMV did not differ across tasks (both regions: *F*_(2,60)_ ≤ 3.07, *p≥* 0.054). PreSMA showed a main effect of task (*F*_(2,60)_ = 4.24, *p* = 0.019, *ɛ*= 0.84), but activation during production did not differ from either perception (*t*_(30)_ = −0.84, *p* = 0.406, *d* = −0.09) or NTFD (*t*_(30)_ = 2.01, *p* = 0.054, *d* = 0.18). Only the SMA showed a strong main effect of task (*F*_(2,60)_ = 24.63, *p* = 1.55×10^−8^, *ɛ* = 0.89), with higher activity for production than for perception (*t*_(30)_ = 6.27, *p* = 6.51×10^−7^, *d* = 0.83) and NTFD (*t*_(30)_ = 2.74, *p* = 0.010, *d* = 0.31). Its activity was also lower during perception than NTFD (*t*_(30)_ = −5.05, *p* = 2.03 × 10^−5^, *d* = −0.51), giving a graded profile across tasks (production > NTFD > perception) rather than a preference for production alone. In the dorsal striatum, by contrast, perception and NTFD did not differ (*t*_(30)_ = −1.11, *p* = 0.276, *d* = −0.16), so only production stood apart from the other two tasks. A two-way RM ANOVA comparing the task profiles of the dorsal striatum and the SMA confirmed this contrast with a significant ROI × Task interaction (*F*_(2,60)_ = 17.94, *p* = 7.82 × 10^−7^, *ɛ* = 0.84).

A similar preference for the production task was observed in the cerebellum, reflected in a significant main effect of task (*F*_(2,60)_ = 11.51, *p* = 5.86 × 10^−5^, *ɛ* = 0.91), with no evidence that task-related differences depended on sensory modality (Modality × Task: *F*_(2,60)_ = 0.56, *p* = 0.575, *ɛ* = 0.87). Post hoc comparisons confirmed higher encoding-related activity for production relative to both perception (*t*_(30)_ = 3.15, *p* = 0.004, *d* = 0.26) and NTFD (*t*_(30)_ = 4.19, *p* = 2.27 × 10^−4^, *d* = 0.38). However, the magnitude of these effects was markedly smaller than that observed in the dorsal striatum, and the cerebellar activation profile remained functionally closer to the cortical cluster in the MDS space (Extended Data Fig. E3b and c). Consistent with this organization, subject-wise correlations with premotor regions were significantly higher for the cerebellum than for the dorsal striatum (all: *t*_(30)_ *≥* 2.15, *p ≤* 0.039, *d ≥* 0.51). This difference could not be attributed to lower reliability in the dorsal striatum, as its *Spearman–Brown* estimate exceeded that of the cerebellum (0.60 *vs.* 0.54; see Methods).

Together, these results reveal a largely modality-invariant cortical core, unlike the sensory cortices, with the cerebellum clustering with PreSMA and SMA. The dorsal striatum stood apart through its task-specific modulation, most strongly preferring the production task.

### Auditory Beat-Based Temporal Prediction Selectively Engages the Dorsal Striatum

With this shared architecture in place, we anticipated greater striatal activity for beat-based sequences and greater cerebellar activity for interval-based ones.

We tested this prediction using a two-way RM ANOVA on the PSC averaged across tasks, with ROI (dorsal striatum, cerebellum) and temporal structure (beat, interval) as within-subject factors. For the auditory modality, the ANOVA revealed the expected interaction between the two factors (*F*_(1,30)_ = 16.61, *p*= 3.11 × 10^−4^, *ɛ* = 1; Figure 5). Post hoc comparisons probing effects of temporal structure within each ROI (Holm– Bonferroni: two ROI-specific tests, *α*_FWE_ = 0.05, smallest *p ≤* 0.05*/*2) showed that PSC was significantly higher for auditory beat than interval sequences in the dorsal striatum (*t*_(30)_ = 3.08, *p*= 0.004, *d*= 0.15), whereas the cerebellum showed no significant difference between them (*t*_(30)_ = −0.78, *p* = 0.440, *d* = −0.04).

**Figure 5.**
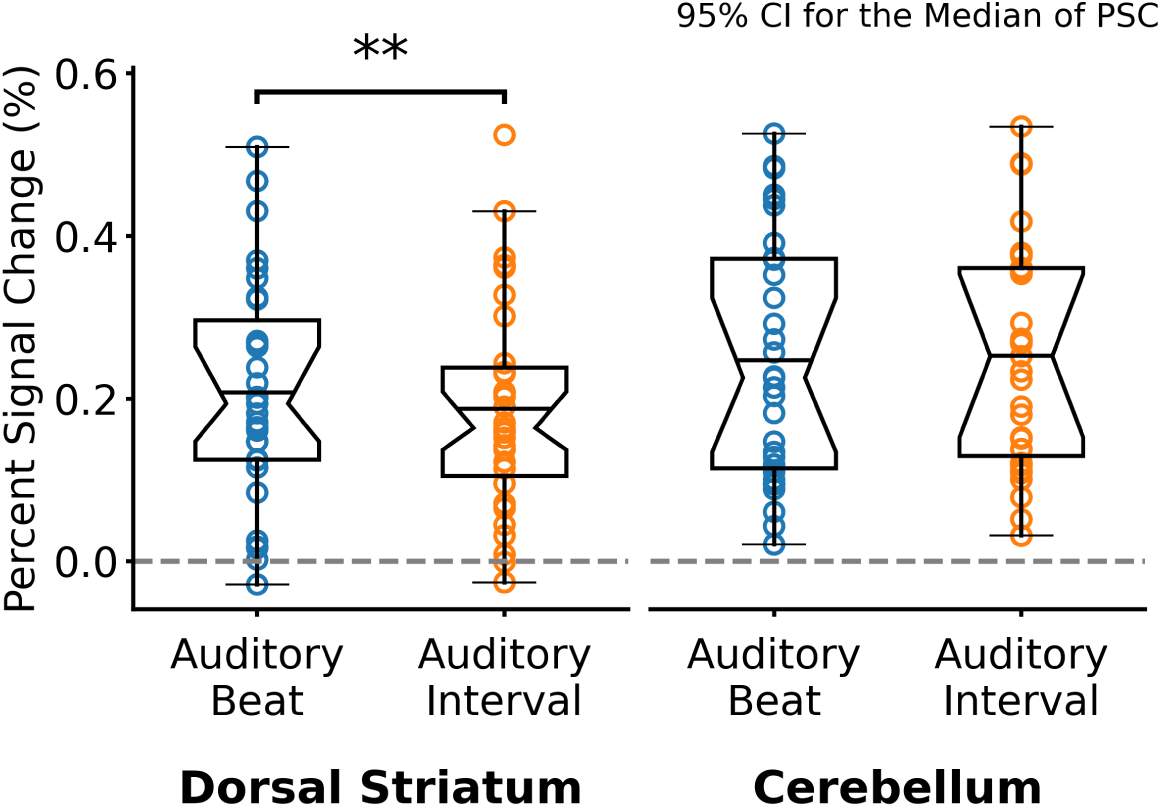
Single dissociation between dorsal striatum and cerebellum during auditory beat-based temporal encoding. A two-way RM ANOVA (ROI × Temporal Structure) reveals beat > interval in the dorsal striatum but not in the cerebellum for the auditory modality. Boxplots show PSC during encoding relative to rest for the auditory beat and interval conditions, averaged across tasks. Boxes represent the IQR, notches the bootstrap confidence interval of the median, whiskers extend to 1.5×IQR, and horizontal lines denote the mean. The annotation indicates a significant post hoc comparison (** 0.001 *< p <* 0.01). The dashed horizontal lines mark PSC = 0.

Task by task, the interaction held in NTFD (*F*_(1,30)_ = 6.54, *p* = 0.016, *ɛ* = 1) and in production (*F*_(1,30)_ = 10.60, *p* = 0.003, *ɛ* = 1), but not in perception (*F*_(1,30)_ = 1.44, *p* = 0.240, *ɛ* = 1). The dorsal-striatal preference for the beat reached significance on its own only in NTFD (*t*_(30)_ = 2.60, *p* = 0.014, *d* = 0.23); it pointed the same way in production (*t*_(30)_ = 1.65, *p* = 0.110, *d* = 0.10) and was absent in perception (*t*_(30)_ = 0.47, *p* = 0.645, *d* = 0.05). Within a single task, the dorsal-striatal difference between beat and interval was thus significant only in NTFD, and it was largest with tasks pooled.

For the visual modality, the same two-way RM ANOVA revealed no significant interaction between ROI and temporal structure (*F*_(1,30)_ = 0.81, *p* = 0.376, *ɛ* = 1), and no significant main effect of temporal structure was observed (*F*_(1,30)_ = 2.75, *p* = 0.108, *ɛ* = 1). These results indicated no differential response to regularity in temporal structure for the visual modality.

The contrasting results across modalities point to auditory-specific beat selectivity in the dorsal striatum. To formally test this, we conducted a two-way RM ANOVA on the dorsal-striatal PSC with modality (auditory, visual) and temporal structure (beat, interval) as within-subject factors. This analysis revealed a significant interaction between modality and temporal structure (*F*_(1,30)_ = 9.84, *p* = 0.004, *ɛ* = 1), where beat selectivity was absent in the visual modality (*t*_(30)_ = −1.24, *p*= 0.225, *d*= −0.07), so the dorsal-striatal difference between beat and interval was present in the auditory modality only. With modalities pooled, the ROI × Temporal Structure interaction remained significant (*F*_(1,30)_ = 11.60, *p* = 0.002, *ɛ* = 1), but the beat and interval conditions no longer differed within the dorsal striatum (*t*_(30)_ = 1.20, *p* = 0.238, *d* = 0.05) or the cerebellum (*t*_(30)_ = −1.63, *p* = 0.113, *d* = −0.08).

The dorsal-striatal preference for auditory beat over interval sequences could reflect a difference already present in sensory processing, or one shared with the rest of the architecture. We therefore submitted the encoding-period PSC of every other ROI, measured in the runs of the main tasks, to the same two-way RM ANOVA, with modality and temporal structure as within-subject factors. The auditory cortex showed a main effect of modality (*F*_(1,30)_ = 310.51, *p* = 2.26 × 10^−17^, *ɛ* = 1), with neither an effect of temporal structure (*F*_(1,30)_ = 0.56, *p*= 0.459, *ɛ*= 1) nor a Modality × Temporal Structure interaction (*F*_(1,30)_ = 0.24, *p* = 0.628, *ɛ* = 1). In no other region did temporal structure produce a main effect or interact with modality (all *F*_(1,30)_ ≤ 2.66, *p ≥* 0.113). Only the dorsal striatum therefore showed a difference between beat and interval. Together, the dorsal striatum, but not the cerebellum, was selectively engaged by auditory periodic sequences; with modalities pooled, this striatal effect did not hold. Yet both sequence types were at least partly predictable, leaving open whether the rest of the architecture depends on predictability or responds to any sequence.

### Temporal Unpredictability Increases Cortico–Cerebellar Activity but Not Dorsal-Striatal or SMA Activity

Apart from the dorsal striatum, no region showed a difference between beat and interval. Yet both were at least partly predictable: beat onsets fell on a periodic pulse, whereas in the interval condition only the final event’s timing could be anticipated, from the standard presented twice before it. These regions may therefore require at least partly predictable timing, or respond to any temporal sequence. To decide between them, we acquired two further NTFD runs, adding a random condition to the beat and interval conditions. Random sequences preserved the trial structure but removed all temporal regularity from the encoding events (there was no repeated standard). All analyses here draw on these runs alone, so their beat and interval trials are a separate acquisition from the main runs reported above. We used two measures: the mean response across nonrandom (beat and interval pooled) and random sequences relative to rest, to test whether a region responds to a temporal sequence regardless of its structure; and the difference between random and non-random sequences, to test whether a region’s response depends on temporal regularity.

The first measure, i.e. the mean response of the temporal-encoding regions defined from the encoding period of the main tasks, was above rest in every region (one-sample *t*-tests against zero, Holm–Bonferroni-corrected across the eight ROIs; dorsal striatum: *t*_(30)_ = 2.35, *p* = 0.025, *d* = 0.42; all others: *t*_(30)_ *≥* 7.37, *p ≤* 3.31×10^−8^, *d ≥* 1.32; Figure 6a). For the second measure, the encoding-period PSC of these runs was submitted to a twoway RM ANOVA with modality (auditory, visual) and predictability (non-random, random) as within-subject factors. In the sensory cortices, activity differed between modalities, and only the auditory cortex also differed between non-random and random sequences: a main effect of modality (*F*_(1,30)_ = 225.66, *p* = 1.69 × 10^−15^, *ɛ* = 1), a main effect of predictability (*F*_(1,30)_ = 14.61, *p* = 6.21 × 10^−4^, *ɛ* = 1), and a Modality × Predictability interaction (*F*_(1,30)_ = 12.98, *p* = 0.001, *ɛ* = 1), with post hoc comparisons (Holm–Bonferroni: two modalityspecific tests, *α*_FWE_ = 0.05, smallest *p ≤* 0.05*/*2) showing lower activity for non-random than random sequences in audition (*t*_(30)_ = −5.77, *p* = 2.62 × 10^−6^, *d* = −0.44) but not in vision (*t*_(30)_ = −0.25, *p* = 0.803, *d* = −0.04). The visual cortex showed only a main effect of modality (*F*_(1,30)_ = 68.74, *p* = 2.97 × 10^−9^, *ɛ* = 1), with neither an effect of predictability (*F*_(1,30)_ = 2.65, *p* = 0.114, *ɛ* = 1) nor a Modality × Predictability interaction (*F*_(1,30)_ = 1.16, *p* = 0.290, *ɛ* = 1). Beyond the sensory cortices, PMD (*F*_(1,30)_ = 15.87, *p* = 4.00 × 10^−4^, *ɛ* = 1), PMV (*F*_(1,30)_ = 15.34, *p* = 4.80 × 10^−4^, *ɛ* = 1), PreSMA (*F*_(1,30)_ = 12.74, *p* = 1.23 × 10^−3^, *ɛ* = 1) and the cerebellum (*F*_(1,30)_ = 6.75, *p* = 0.014, *ɛ* = 1) showed a main effect of predictability, with greater activity for random sequences, and neither a main effect of modality nor a Modality × Predictability interaction in any of them (all: *F*_(1,30)_ ≤ 0.79, *p ≥* 0.381). Thus, activity in PMD, PMV, PreSMA and the cerebellum was greater for random sequences in both modalities, and in the auditory cortex for auditory sequences only, whereas activity in the visual cortex differed only between modalities.

**Figure 6.**
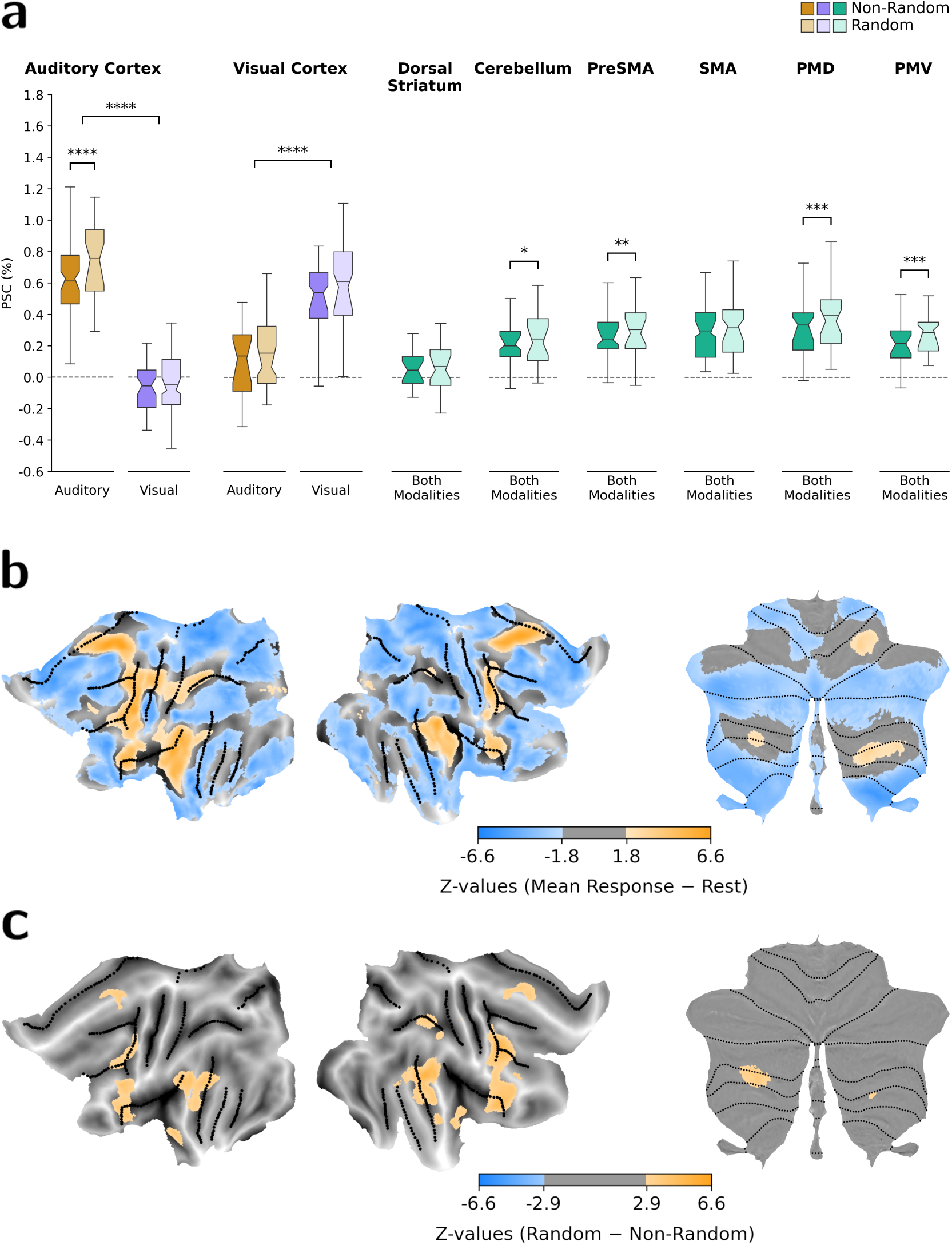
Cortico–striatal–cerebellar profiles of temporal unpredictability. **(a)** PSC relative to rest for non-random (the average of beat and interval) and random sequences, assessed with two-way RM ANOVAs (Modality [auditory, visual] × Predictability [non-random, random]). Annotations mark effects surviving omnibus and, where applicable, FWE-corrected post hoc testing (*α*_FWE_ = 0.05; * *p <* 0.05, ** *p <* 0.01, *** *p <* 0.001, **** *p <* 10^−4^). Both-modalities plots reflect a main effect of predictability only; modality-specific plots reflect a main effect of modality (bracket spanning modalities) and, where present, a Modality × Predictability interaction (within-modality annotation). Boxes show the IQR, notches the bootstrap 95% CI of the median, whiskers extend to 1.5×IQR, and horizontal lines denote the mean; dashed lines mark PSC = 0. **(b)** Mean encoding response across non-random and random sequences, relative to rest: regions responding to temporal sequences regardless of their structure (orange: sequences > rest; blue: rest > sequences). **(c)** Difference between random and non-random sequences: regions whose response differs between random and non-random sequences (orange: random > non-random; blue: non-random > random). Random sequences produced greater activity than non-random sequences, and almost no region showed the reverse effect. In **(b)** and **(c)**, group *z*-maps are shown on cortical (fs LR32k) and cerebellar (SUIT) flatmaps, thresholded at a two-sided whole-brain FDR criterion (*q ≤* 0.05) computed per map.

In contrast, the dorsal striatum and the SMA showed no effect of predictability (dorsal striatum: *F*_(1,30)_ = 3.07, *p* = 0.090, *ɛ* = 1; SMA: *F*_(1,30)_ = 1.66, *p* = 0.207, *ɛ* = 1), nor any main effect of modality or Modality × Predictability interaction (dorsal striatum: both *F*_(1,30)_ ≤ 0.05, *p ≥* 0.826; SMA: both *F*_(1,30)_ ≤ 0.16, *p ≥* 0.695; Figure 6a).

Both measures were then mapped across the whole brain. The mean response was positive throughout the temporal-encoding regions and negative across the *Default-Mode Network* (Figure 6b). Significant differences between random and non-random sequences lay within those same regions, in portions of the premotor cortex (PreSMA, PMD, and PMV), the posterior auditory cortex extending into the Supramarginal Gyrus, and the inferior *Demand* subdivision of the cerebellum [28], and almost all reflected greater activity for random sequences (Figure 6c and Extended Data Table E2). Outside these regions, only the anterior insula showed a difference: its mean response did not differ from rest, and its activity rose above rest for random sequences only (Supplementary Note 1).

The temporal-encoding regions therefore respond to temporal sequences regardless of their structure, and most respond more strongly when the sequence is irregular. The dorsal striatum and the SMA are the exceptions: their activity did not differ between regular and irregular sequences.

## Discussion

We asked whether the division of labour between the basal ganglia and the cerebellum holds across sensory modalities and tasks. Temporal encoding recruited a similar cortico–striatal–cerebellar architecture in every combination of temporal structure, sensory modality and task. Two properties set the dorsal striatum apart: a selectivity for the production task that no cortical region matched, and a greater response to the auditory beat than to the interval condition, which the cerebellum did not show. This response was selective for the beat, not for regularity: removing regularity altogether, in random sequences, did not lower it.

The subcortex has been proposed to represent time, and the cortex the task goal [3]. This proposal rests on two findings. First, explicit and implicit timing recruit non-overlapping cortical areas [12, 31]. Second, both show the same cerebellar and basal-ganglia dissociation for regularity, whether timing is explicit [5, 6] or implicit [3]. Our data do not support this. With encoding matched across tasks and only the response varying, the same premotor and cerebellar regions were engaged whether the duration was reported explicitly (production, perception) or used implicitly (NTFD). Task goals did modulate the system, but most strongly outside the cortex: the dorsal striatum showed the largest task-related effects of any region examined.

These task effects follow how prospectively each task uses the temporal estimate: how far ahead of the final timing event it is needed. This is greatest in production, where the estimate times the movement; intermediate in NTFD, where it prepares for the final to-be-judged event; and least in perception, where it is evaluated only once that event has passed. The dorsal striatum and the cerebellum responded more only at the top of this scale, in production, with a far larger effect in the dorsal striatum (*d ≥* 1.10 *vs. d ≤* 0.38). What sets production apart is not its motor output, since all three tasks end in a button press, but that the estimate determines when the movement begins, the operation for which the dorsal striatum is classically required [4, 31, 32]. Consistent with this, the striatal encoding peaks fell in the executive subdivision, defined by its structural connectivity with the dorsolateral prefrontal cortex [30]. The cerebellar peaks fell in the *Demand* regions, functionally connected with the dorsal attention and control networks, and the clusters extended into the *Motor* regions, connected with the somatomotor and premotor networks [28]. The SMA—the only cortical region with a strong task effect—instead showed a graded decrease in activity from production to NTFD to perception, mirroring this scale, as expected from its greater engagement in motor than perceptual timing [11]. Through its loops with the dorsal striatum and the cerebellum [13], it may pass its graded signal to both, which respond only when the estimate starts a movement.

Our single dissociation does not contradict the double dissociation on which the beat–interval division rests [3]. In that study, healthy participants benefited from single-interval and rhythmic cues to a similar degree relative to random sequences (Cohen’s *d* between 1.4 and 1.8 in two control groups), so neither form of prediction was privileged in the intact brain. The dissociation appeared only under damage, which tests whether a structure is necessary: with cerebellar damage, the single-interval benefit was lost while the rhythmic benefit remained large (*d* = 0.92); with the basal ganglia compromised, the pattern reversed. Their rhythmic and single-interval cues also differed in more than periodicity: the rhythmic cue presented the target duration three times, immediately before the target, whereas the single-interval cue presented it once, followed by a variable delay. In our study, the standard was presented equally often in the beat and interval conditions, and the delay before the response was matched on average, so differences between them can be attributed to periodicity. Assessing encoding activity also asks a graded question: how strongly each structure is recruited while both are intact. Both responded in every condition (Figure 4), and only the dorsal striatum responded more to the auditory beat than to the interval. A single dissociation is therefore compatible with the lesion evidence: a structure can be necessary for one form of prediction without responding more strongly to it.

This pattern allows two interpretations: the dorsal striatum amplifies a timing signal already encoded by the cerebellum, received through disynaptic cerebello-striatal connections [13], or it acts independently of that signal, as a default beat-based timer [7]. Under the first, beat-based prediction should fail when the cerebellum degenerates, leaving no signal to amplify; yet the rhythmic benefit is preserved under cerebellar degeneration [3]. The dorsal striatum therefore does more than amplify a cerebellar timing signal: it can sustain periodic prediction on its own. Under the second, the cerebellum would instead correct timing errors, which fits our observation of equal responses to the beat and the interval and a greater response to random sequences. It may therefore still contribute to periodic prediction. The patient data cannot settle this: the rhythmic benefit under cerebellar degeneration did not differ from controls, which shows that the cerebellum is not necessary for it, but does not rule out a smaller contribution. Compensation, already argued against [3], could take two forms: either the patients acquired a beat-specific striatal signal after damage, or the damage strengthened a signal that was already present. Our results rule out the first, since the signal is present in brains that were never damaged; the second remains to be tested.

The double dissociation in patients was obtained with visual sequences [3], whereas our striatal effect was auditory: the visual beat and interval conditions did not differ, and the beat effect was reliably larger in audition than in vision. Their rhythmic sequences were also isochronous, whereas our beat was implied by integerratio spacing. This choice has a cost: a sparse, implied beat is likely a weaker inducer than an isochronous stream [16], so effects reported with isochronous sequences may be smaller or absent here. The measures also differed: theirs asked whether behaviour breaks down under damage [15], ours whether encoding activity differs between temporal structures. Visual sequences did engage the dorsal striatum, but their periodicity did not increase that engagement, possibly because an implied beat is harder to induce visually: a regular visual rhythm engages the basal ganglia once audition has primed a beat in it [33]. Since striatal activity reflects the prediction of an established beat rather than the search for one [10], visual sequences that induced no beat would give the dorsal striatum nothing to predict.

We used random sequences to test whether the striatal response to the beat condition reflects a preference for predictability. If so, striatal activity should have been lowest when regularity was absent; instead, it was no lower for random than for non-random sequences. The response therefore appears to be driven by the presence of the beat, not by the predictability that the repeated standards also gave the interval condition. The contrast with random sequences could only be tested within NTFD, a task in which the dorsal striatum also showed the beat preference. In other regions, activity increased when the stream became irregular: reliably in PreSMA, PMD and PMV, and to a lesser extent in the cerebellum. The SMA showed no such increase, and thus patterned with the dorsal striatum, with which it forms a loop [13]. Previous work found the same pattern: rhythms with a beat increased putamen activity and its functional connectivity with the SMA, whereas irregular rhythms increased activity in PreSMA, premotor cortex and the cerebellum [34]. The beat and irregularity thus engage different regions, and the division does not follow the boundary between cortex and subcortex: the dorsal striatum and the SMA fall on the side of the beat, and PreSMA, lateral premotor cortex and the cerebellum on the side of irregularity. Only the anterior insula responded outside this architecture, rising above rest for random sequences only.

Under group-defined regions, the striatal beat effect no longer survived correction (Supplementary Note 2): averaging regions across participants blurs their anatomical and functional variability and dilutes effects in small structures [23]. Group-level studies may therefore have underestimated striatal contributions to beatbased prediction. Importantly, this gain in sensitivity did not come at the cost of bias: our localizing contrast was orthogonal to all tested effects (Methods).

In conclusion, the beat–interval division is real, but not symmetric. The cerebellum was engaged whenever a sequence had to be tracked, and more strongly when no regularity could be exploited. The dorsal striatum was engaged in addition, when the sequence carried an auditory beat and when the estimate had to drive an action. This asymmetry is invisible to damage, which reports what is lost rather than what is at work: hence a double dissociation in patients beside a single dissociation in the healthy brain. Read together, they describe an architecture in which the cerebellum always runs, and the dorsal striatum joins it for the beat, and for the action.

## Methods

To avoid ambiguity with terms related to MRI, we adhered to the definitions provided in the Brain-Imaging-Data-Structure (BIDS) Specification version 1.10.0 [35].

### Participants

Thirty-one adults (16 females), aged 18–41 years at recruitment (mean *±* SD = 24.71 *±* 6.37 years), participated in the study. Participants responded to poster advertisements and were not recruited for any particular expertise, musical or otherwise. They were excluded if they had a history of brain surgery, neurological or psychiatric disorders, used psychotropic medications during the study, or did not meet the safety guidelines of the *Centre for Functional and Metabolic Mapping* (CFMM) at Western’s Robarts Research Institute, London, Ontario, Canada. These guidelines included general MRI safety measures, such as the absence of metal or other ferromagnetic devices in the body and no history of claustrophobia. All participants were compensated a total of 175 CAD for behavioural training and imaging sessions.

### Tasks

Three duration-based timing tasks were designed to probe, through functional imaging, the temporal encoding of sequences that varied in their temporal structure and response requirements.

Three tasks were used: (1) *production*; (2) *perception*; and (3) *Non-Temporal Feature Discrimination* (NTFD). All of them used the same type of encoding and differed only in the response required after it. During the encoding period, events were presented in one of two temporal structures: *beat* or *interval*. Encoding in both cases contained two presentations of a *standard* duration, each followed by a longer, intervening duration.

Both were *Stimulus-Onset Asynchronies* (SOAs), the time between the onsets of successive events; we refer to the longer ones as intervening SOAs. For the beat condition, each of the two intervening SOAs was three times the standard; for the interval condition, the intervening SOAs were jittered (hereafter, jittered SOAs), but their sum equalled six times the standard. Thus, the beat encoding structure contained an underlying periodicity: each intervening SOA was the time it would take for three additional events to sound if the sequence was isochronous. In other words, the beat encoding was composed of eight standard durations, but only the first and fourth durations were marked by stimuli. Because humans generally perceive a beat at SOAs ranging from 450 to 700 ms ([36], chapter 2), we generated five evenly spaced standards within this range: 459 ms, 510 ms, 561 ms, 612 ms, and 663 ms. We selected these values as they were multiples of 17 ms, ensuring compatibility with the 60 Hz screen refresh rate. In all tasks, each trial corresponded to one of the five standards shown in Figure 1, with trial length varying accordingly. Each standard had an equal number of trials for beat and interval within a task. Furthermore, the average of each intervening SOA, and thus the average length of the entire trial, was the same for beat and interval. Additionally, the first jittered SOA in the interval condition was generated by multiplying all standards by three, creating a range of integers between the resulting minimum and maximum values, retaining only those divisible by 16 or 17 to ensure synchronization with the screen refresh rate, and randomly selecting from this set. The second jittered SOA was calculated as six times the standard of the trial minus the first jittered SOA.

Tasks were also presented in auditory and visual modalities. In the auditory tasks, events were *∼*80-ms beeps, whereas in the visual tasks, they were geometric shapes displayed for *∼*80 ms.

Here, *experimental condition* refers to either the encoding or response period within a trial. During the encoding period, conditions corresponded to specific combinations of task, sensory modality, and temporal structure, whereas during the response period they were defined solely by the task. For details about how experimental conditions were modelled as regressors-of-interest, consult Methods.

Each run comprised 30 trials of the same task presented within the same modality (for details on the trial structure and duration of each task condition, refer to Methods). There were 15 trials each for each temporal structure, i.e. beat and interval, in which trials for every standard were repeated three times. Trials were separated by two-second rest periods. In addition, there were also five 6-s trials of rest, experienced by the participant as an extended inter-trial period. For each task within a run, the order of trials—comprising the two temporal structures and rest—was pseudo-randomized across runs, sessions, and subjects. The randomization process ensured that a rest period was not presented as the first or last trial, did not appear consecutively, and did not occur at every other trial.

Stimuli were displayed on a grey screen with a black fixation cross at the centre. The fixation cross remained visible, except during presentation of the shapes in the visual task.

The behavioural protocols for each task, along with the generation of SOAs and their randomization across trials, are openly available as outlined in Methods.

### Production Task

The production task examined how accurately the participant reproduced the timing of the standard duration for beat and interval sequences. On each trial, participants were presented with two standard durations separated by two intervening SOAs, and then a final event that marked the beginning of a third standard duration. Participants then pressed a button with their index finger at the time they judged to accurately reproduce the end of the standard duration. When they pressed the button, a corresponding event (beep or flash, depending on the modality) occurred to provide feedback. Participants had two seconds to respond (see Figure 1).

Trial durations ranged from 5752 ms (shortest standard) to 7384 ms (longest standard), and a complete run—consisting of the production task completed in each modality—lasted 10 minutes and 4 seconds.

### Perception Task

The perception task examined how accurately the participant judged whether a deviant final duration was shorter or longer than the standard durations, for beat and interval sequences. On each trial, participants were presented with two standard durations separated by two intervening SOAs, and then a final *comparison* duration that was either longer or shorter than the standard. Concretely, these comparisons were 2%, 12%, or 20% longer or shorter than the standard. Trials featuring a specific comparison for a given standard were evenly distributed across the two imaging sessions, ensuring that each modality included one instance of every standard-comparison pair within each session. Afterwards, a panel was displayed prompting participants to respond whether the comparison was longer or shorter than the standard. Participants pressed the button under their index or middle finger, respectively, within a fixed two-second response period.

Trial duration ranged from 6120 ms (shortest standard and comparison) to 8180 ms (longest standard and comparison), and a complete run—consisting of the perception task completed in each modality—lasted 10 minutes and 38 seconds.

### Non-Temporal Feature Discrimination Tasks (NTFD)

The *Non-Temporal Feature Discrimination* (NTFD) task examined how quickly participants completed a simple, non-temporal stimulus discrimination during beat and interval sequences. In each trial, participants were presented with two standard durations separated by two intervening SOAs, and then a final standard duration, with the final event changed from those prior. In the auditory modality, the final event was either higher (880 Hz) or lower (220 Hz) than the preceding events (440 Hz). In the visual modality, the final event was either a circle or a triangle, while all preceding events were rectangles. Participants were instructed to promptly identify the final event as either higher/lower or circle/triangle, by pressing the button under their index or middle finger, respectively. Participants had 2000 ms to respond. Each button press produced an event (beep or flash, according to the modality) as feedback (see Figure 1).

Trial duration ranged from 6211 ms (shortest standard) to 8047 ms (longest standard). A complete run— consisting of the NTFD task completed in each modality—lasted 10 minutes and 38 seconds. For each modality, trials per temporal structure and discrimination-response type were balanced across runs within a session. Since the beat and interval conditions each consisted of 15 trials per task, if the beat condition included eight trials with a higher beep response in the first auditory task of the session, it would include seven such trials in the second task, and vice versa.

In a variation of the NTFD task, we introduced a third trial type, *random*, in distinct runs, in which the sequence timing during encoding was entirely unpredictable. In these runs, participants completed 10 trials for each of the three temporal structures (beat, interval, and random), with five trials assigned to each discrimination response. We pseudo-randomized the durations between events, ensuring that no two consecutive durations were shorter than the original shortest standard (459 ms) and that all durations were multiples of either 16 ms or 17 ms to match the 60 Hz screen refresh rate. We defined five types of random trials and matched the total trial duration for each type to a corresponding beat or interval trial. As a result, these runs had the same total duration as the runs featuring only beat and interval trials.

### Behavioural Training

Three behavioural training sessions, each approximately 3 hours in duration, were conducted in the testing rooms at the Western Interdisciplinary Research Building of Western University. These sessions ensured participants were comfortable and proficient with the experimental tasks, reducing variability due to learning effects and individual differences.

In the first session, participants received general instructions about the execution and timing of all tasks. Concretely, written instructions were provided for each task. Then, a brief demonstration and practice preceded the execution of every single task.

Across all three sessions, participants completed the tasks on a laptop, which presented both visual and auditory stimuli, and responded according to the provided instructions. We note that training and imaging sessions were identical in terms of presentation of stimuli and execution of tasks.

Preventing saccadic movements and keeping perimetric origin of the visual field constant during imaging data acquisition, as means to avoid spurious effects in the functional maps, were critically important. Training sessions accounted for this predicament and, thus, participants were asked to move their eyes as little as possible while fixating towards a cross placed at the centre of the screen. In addition, participants were instructed to provide their button-press responses using the laptop keyboard that was utilized to display the stimuli. They used their right hand, with the same fingers assigned to each response type, as if operating the MRI-compatible response box (see Methods for details).

The first session consisted of 24 uninterrupted runs, organized into four loops. Each run, lasting approximately three and a half minutes, was dedicated to a single task. Within each loop, all tasks were presented once in a randomized order for each participant, resulting in four runs per task throughout the session. Since the three sessions for each participant were conducted a few days apart, no initial debriefing was performed during the second and third sessions, allowing for more runs to be included. These remaining two sessions thus consisted of 26 runs, with four additional runs dedicated to the *perception* tasks. We note that in these training sessions each run was shorter than its corresponding run in the imaging sessions, because: (1) the response period ended immediately after a response was made; (2) duration between trials was shorter, i.e. 1.504 s instead of 2 s; and (3) no baseline period was included. For details of the experimental designs of the tasks, consult Methods and Figure 1.

All thirty-one participants, who completed both behavioural training and the imaging sessions, reported that they had understood all tasks before going into the scanner. The experimenters did not observe, and the participants did not report, any significant difficulties in executing the tasks by the end of the third training session. To verify that participants followed the instructions and performed the tasks correctly, confirmatory analyses of the behavioural data from the three sessions were conducted prior to proceeding with the imaging data collection (see Methods for more details).

### Imaging Sessions

Each participant completed two imaging sessions, with each session approximately two hours long.

Upon arriving at CFMM, participants were rebriefed on task execution. This short training was conducted in a designated room outside the scanner environment. In the first session, participants also completed a screening form that covered safety procedures, general health, and medical history prior to entering the scanner. Participants were scanned in the supine position and were instructed to minimize head movement. Foam pads were positioned within the coil for comfort and stability. Motor responses were recorded using a fourbutton response box operated with the right hand (see Methods for details). Earplugs were provided to reduce scanner noise, and headphones were used to enable verbal communication with the experimenters in the control room.

Each functional MRI run involved the alternating presentation of a task for each modality, ensuring thus that each task paradigm was presented twice per session (consult Methods for detailed information about the tasks). After the initial Transistor-Transistor Logic (TTL) pulse at the start of the run, a twelve-second rest period was introduced prior to the onset of the first task to ensure stabilization of the *Blood-Oxygen-LevelDependent* (BOLD) signal. Each run also included a six-second rest period between tasks and a ten-second rest period at the end. Data from each of the six tasks were collected twice across two separate runs within the same session. The two sessions included all main tasks, with the second session featuring an additional variation of the NTFD task. Specifically, the first session included six fMRI runs for the main tasks plus one anatomical run, while the second session also featured six fMRI runs for the main tasks, along with two additional runs for the NTFD-task variation. The order of the fMRI runs was randomized across sessions and subjects. In addition, both sessions included a gradient-echo run for spatial-distortion correction (for more details about imaging-data sequences, refer to Methods).

The runs were interspersed with brief pauses. During these breaks, experimenters monitored participant status, and visually inspected the newly acquired MRI data for quality assurance.

### Questionnaires

At the beginning of the first training session, participants completed a demographic questionnaire. Postexperiment questionnaires were also completed at the end of each training and imaging session to collect feedback on their overall experience.

### Stimuli

The stimuli were presented using custom scripts programmed in Python 3.7.11 with Expyriment v0.10.0 [37]. Because of observed timing errors, a second version of the auditory protocols was subsequently implemented in PsychoPy v2022.2.5 [38] and used to collect a second behavioural cohort on the auditory tasks only. The two versions used identical stimuli and the same audio interface, and differed only in the presentation software.

Auditory stimuli were delivered through a *Steinberg* UR22mkII 2×2 USB 2.0 Audio Interface. Sine-wave tones at 220 Hz, 440 Hz, and 880 Hz, each 80 ms in duration, were generated in MATLAB R2020b.

To verify stimulus timing, we retrospectively recorded stimulus onsets with a Cedrus StimTracker Duo. Visual onsets were detected with a photodiode mounted on the display, and auditory onsets were recorded from the signal relayed by the Steinberg UR22mkII interface, each event being time-stamped independently of the presentation computer. We replayed the sequences offline, with no participant, to measure their rendered onset timing. In total, we recorded the 360 sequences of two participants’ stimulus sets, spanning all modalities, tasks, and conditions, with each sequence recorded two to eight times (1290 recordings). Comparison of rendered and intended onset timing revealed timing variability in the stimulus presentation, which was greater for auditory than visual stimuli.

To test whether timing errors were reproducible across presentations, we computed one-way *Intraclass Correlation Coefficients* (ICCs) of the per-duration timing error (rendered minus nominal duration) across repetitions of the same sequence. The ICCs were near zero (−0.01 auditory, 0.05 visual), indicating errors were not reliably associated with particular sequences. The variance of the difference between two recordings of a sequence was about twice the error variance of one recording (2.03 auditory, 1.95 visual), close to the value of 2 expected when errors are independent across recordings. Thus, the timing errors were not sufficiently reproducible to identify and exclude a stable subset of bad trials.

We characterized timing fidelity across conditions, starting with the auditory beat condition, the one in which regular timing supports beat induction. The pulse of a beat sequence is set by the structure of its durations: within each sequence, the standard *S* is marked by two tones, and the intervening SOA between pairs of tones is 3*S*, so every duration is an integer multiple of *S*, the structure that induces a beat [17]. For each rendered duration *d_i_* we calculated its deviation from the corresponding nominal duration *n_i_* and summarized the timing errors using the *Root-Mean-Square* (RMS) deviation in milliseconds, 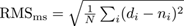 and as a percentage of each duration’s nominal value, 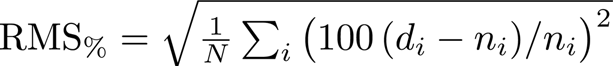. Normalizing each error by its own nominal duration allowed us to characterize relative timing variability across durations of different lengths. In the auditory beat condition, the RMS deviation was 2.66% (15.5 ms) for the standards and 1.20% (19.6 ms) for the intervening SOAs. The RMS, however, summarizes typical deviations and conceals the larger ones. We therefore also counted the trials on which a rendered duration exceeded two published thresholds: the Weber fraction for duration discrimination of empty intervals, about 5% [39], and the timing variability of 8.6% at which pulse extraction was reported to cease in a fully sounded sequence [40]. No intervening SOA exceeded either threshold, but a standard deviated by more than 5% in 27.6% of trials and by more than 8.6% in 4.8%. The ratio between an intervening SOA and a standard, nominally 3, deviated by more than 5% in 24.5% of trials and by more than 8.6% in 4.8%, all of them among the trials with a deviating standard (Supplementary Note 3). Which of these deviations disrupted the beat percept cannot be determined, and disruption on some trials may also have discouraged a beat-based strategy on others; the beat percept may therefore have been weakened beyond the affected trials.

In the visual beat condition, the timing variability was smaller, with RMS deviations of 1.46% (7.9 ms) for the standards and 0.42% (6.7 ms) for the intervening SOAs, and no standard, intervening SOA or ratio between them deviated by more than 5% from its nominal on any trial (Supplementary Note 3).

The interval conditions were aperiodic and were designed not to induce a beat. We therefore characterized the timing of the standards, the durations relevant to the participant’s judgements or production responses. They deviated by 2.86% (15.9 ms) in the auditory modality and 1.45% (7.8 ms) in the visual.

Across both temporal structures, the mean signed timing errors were close to zero: 0.00% for the auditory durations and −0.03% for the visual. This indicates the errors did not produce an overall tendency towards lengthening or shortening.

In the perception task, the participant judged a comparison duration against the standard of the same trial. Timing errors could affect these judgements in two ways: a systematic difference between rendered comparisons and standards could shift the point of subjective equality (PSE), whereas trial-to-trial variability could reduce discrimination precision and increase the difference limen (DL). The signed mean deviations of the comparisons were small (within 0.3% for auditory stimuli, 0.15% for visual stimuli), as were those of the standards. Thus, there was no appreciable systematic offset between nominal and rendered durations that would be expected to shift PSE. Timing variability was not zero (RMS *∼* 3.6% auditory, *∼* 1% visual). Under the assumption that these errors were independent and unbiased, they would be expected to add variability to duration judgements, potentially inflating the DL. Both PSE and DL were estimated by the Method of Constant Stimuli (Methods).

For the subsequent PsychoPy re-implementation, which used the same auditory stimuli and interface, we recorded the 90 auditory sequences of a third participant’s stimulus set four times each (360 recordings). The re-implementation rendered them with substantially smaller timing deviations: in the auditory beat condition, RMS deviations were 0.86% (4.8 ms) for the standards and 0.31% (5.1 ms) for the intervening SOAs. Across all auditory sequences, the residual deviations remained random and unbiased (one-way ICC 0.01, variance ratio 1.98). This pattern is consistent with the original timing variability arising from the interaction between the presentation software and audio hardware. Behavioural performance in the second cohort (tested on the PsychoPy re-implementation) did not differ significantly from the imaging cohort on all three tasks. This provides some reassurance that the timing variability did not produce an appreciable change in task performance. However, the absence of a difference does not establish that the jitter had no effect on beat extraction or other behavioural measures (Supplementary Note 4).

Deviations above the two published thresholds occurred on a subset of auditory beat trials, and the jitter may therefore have reduced the temporal regularity of individual beat sequences. Because the distinction between beat and interval was central to the experimental manipulation, this possibility is relevant when interpreting the results. The second implementation substantially reduced timing variability without producing a detectable behavioural difference between cohorts, supporting the robustness of the behavioural findings while not ruling out more subtle effects of timing variability on beat salience. To the extent that the reported results (Results) reflect responses to the implied beat, and that jitter only weakens regularity, the behavioural and imaging beat versus interval results likely reflect lower bounds on the true effects.

The behavioural protocols for both versions are publicly available, as detailed in Additional Information.

### MRI Equipment

The MRI data were acquired using a Siemens 3T Magnetom Prisma Fit MRI scanner with a Siemens 32-channel head coil. A 4-button HHSC 1×4 L Fibre Optic Response Pad was used to record participants’ behavioural responses during the imaging sessions. Sensimetrics S14 Insert Earphones connected to an Avotec Silent Scan 3000 Audio System were used for audio transmission inside the scanner. Visual stimuli were displayed on an MR-compatible screen (Avotec sV-6011 Rear Projection Video, 1024 × 768 resolution) and presented to the participant via a detachable double mirror mounted on the head coil.

All imaging sessions were conducted at the CFMM.

### Data Acquisition

#### Behavioural and Demographic Data

Participants were required to provide overt responses in all tasks. Behavioural data, including response choices (alpha-coded) and stimulus-response latencies (in milliseconds), were recorded by the stimulus-delivery software (refer to Methods for details).

The age and sex of the participants were extracted from the demographic questionnaires (see Methods).

#### Imaging Data

Functional MRI data were collected in runs using a Gradient-Echo (GE) pulse, whole-brain Multiband (MB) accelerated [41, 42] Echo-Planar Imaging (EPI) T2*-weighted sequence with BOLD contrasts, using the following parameters: Repetition Time (TR) = 1200 ms; Echo Time (TE) = 30 ms; flip angle = 40°; Field-of-View (FoV) = 210×210×143 mm^3^; and voxel size = 2.5×2.5×2.5 mm^3^. Slices were acquired in axial orientation, with posterior-to-anterior phase-encoding direction and interleaved sequence. In-plane acquisitions were accelerated by a factor (GRAPPA) of 2; an MB factor of 3 was used across slices.

Gradient Echo Field Maps were acquired in dedicated runs to account for correction of spatial distortions due to B0 inhomogeneities. The following parameters were used: TR = 500 ms; TE1 = 4.92 ms and TE2 = 7.38 ms; flip angle = 60°; FoV = 240 × 240 × 173 mm^3^; and voxel size = 3 × 3 × 3 mm^3^.

A whole brain 3D Magnetization-Prepared RApid GE (MPRAGE) T1-weighted anatomical-image volume was also acquired with the following parameters: voxel size = 1×1×1 mm^3^; flip angle = 9°; and FoV = 256×256×264 mm^3^. Slices were acquired in sagittal orientation.

Additionally, physiological signals for heartbeat and respiration were recorded in real-time during each EPI run.

### Data Analysis

#### Analysis of Behavioural Data

Behavioural performance on each task was summarized by a single perparticipant metric, computed separately for each temporal structure, i.e. beat and interval.

For the production task, we computed the *Signed Asynchrony* (SA), the trial-level deviation between the participant’s *Produced Duration* (PD) and the *standard* (S):

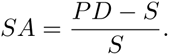

Expressed as a proportion of the standard, SA is the signed relative timing error of the produced duration: positive when the participant over-produces the duration (PD > S), negative when they under-produce it, and zero for a veridical reproduction. The summary metric was the *Mean Signed Asynchrony* (Mean SA): the mean of SA across trials. A non-zero Mean SA thus reflects a *constant error*, i.e. a systematic tendency to overor under-produce the standard.

For the perception task, we summarized discrimination sensitivity by the *Difference Limen* (DL). Under the *Method of Constant Stimuli*, the comparison was presented at fixed proportions of the standard (*±*2%, *±*12%, and *±*20%; Methods), and a logistic psychometric function was fitted by maximum likelihood to the proportion of “longer” responses across comparisons. The DL is half the difference between the two comparison durations judged longer than the standard on 75% and on 25% of trials, expressed as a proportion of the standard. It is the smallest duration change discriminated reliably, so a lower DL denotes finer temporal sensitivity. A DL was estimated for each standard and averaged across standards.

For the NTFD task, we summarized performance by the *Reaction Time* (RT), the latency between the onset of the final (probe) event and the button press, corrected for the fixed presentation and response-device latency of the apparatus. The summary metric was the mean RT across trials.

These three metrics were subsequently compared between the imaging cohort and a second behavioural cohort, to test whether performance changed with the re-implementation of the auditory protocols in PsychoPy (Supplementary Note 4).

For all tasks, bad trials were identified from the log files (Methods) by detecting trials that met predefined exclusion criteria (missing responses, extreme response times, outlier asynchronies, and incorrect discriminations; Methods), and their proportion relative to the total trials per participant was then computed. Confirmatory analyses evaluated each participant’s performance from the behavioural data collected during training. For the production task, a trial was counted as successful when the produced duration deviated from the standard by no more than 25% (i.e. *|PD − S|/S ≤* 0.25), and participants were retained if at least 50% of their trials were successful. For the perception task, participants were retained whose per-standard DL (averaged across conditions and modalities) stayed below the largest comparison (20%) for all five standards. For the NTFD task, participants were retained if their discrimination score, the proportion of correct feature discriminations, was at least 90%.

Descriptive statistics (mean, range, and standard deviation) summarized participants’ age (Methods).

#### Preprocessing of MRI Data

DICOM files were anonymized, converted to NIfTI format, and arranged according to BIDS. This process was facilitated by Autobids (https://github.com/khanlab/autobidsportal), a platform for automated neuroimaging-data standardization that employs Heudiconv (https://github.com/nipy/heudiconv) to convert files and organize data into structured directory layouts.

For data-sharing purposes, facial features were removed from the T1-weighted images with *PyDeface* v2.1.0 [43].

Raw data were preprocessed using the *SPM12* software package [44, 45, Wellcome Department of Imaging Neuroscience, London] v7771. For each subject, an individual coordinate system was first established by positioning the origin of the anatomical image at the approximate location of the anterior commissure.

Voxel displacement maps for GE-EPI volumes of each run were generated using the FieldMap Toolbox v2.1 of SPM12 [46, 47], based on the double-phase and magnitude field map data. Within each session, the GE-EPI volumes were realigned to correct for head motion and head position differences across sessions using a sixparameter rigid body transformation with a second-degree B-spline interpolation [48]. These functional images were subsequently corrected for susceptibility-induced geometric distortions using the voxel displacement maps [49], and resliced together with their mean using a fourth-degree B-spline interpolation. Additionally, the mean functional image was co-registered onto the anatomical image [50], and the resulting transformation was applied to all functional images.

Anatomical images were segmented into grey matter, white matter, *Cerebrospinal Fluid* (CSF), bone, soft tissue, and air/background. The resulting deformation fields were then used for spatial normalization to the standard MNI152 space [51]. The mean functional image and tissue probability maps (grey matter, white matter, and CSF) were used to create a grey matter mask and a whole-brain mask, which were subsequently applied to functional images during post-processing (see Methods).

Neither group normalization nor smoothing was performed at this stage.

For each subject, the corresponding anatomical image was provided as input to FreeSurfer v6.0.0 to generate cortical surface meshes (white-matter and pial tissue interfaces) [52, 53]. These individualized surface meshes were then resampled onto the hemisphere-symmetric fs LR32k template [54, 55] using the *Connectome Workbench* [56].

#### FMRI Model Specification

For each run, corresponding to a specific task paradigm (production, perception or NTFD), the individual fMRI data were analysed using the *General Linear Model* (GLM) as implemented in SPM12. Regressors-of-interest were constructed by convolving the *Haemodynamic Response Function* (HRF) with the temporal profiles of task-specific conditions [57, 58], to capture variations in the BOLD response specific to task performance. The temporal profiles were, in turn, modelled from boxcar functions, specified according to the paradigm descriptors of the tasks. These paradigm descriptors—organized into triplets consisting of onset time, duration, and trial type—were determined from the log-file registries of each subject created by the behavioural protocols.

For each trial, the encoding and response periods were modelled separately. A single regressor was used for the response period across all trials, while four different regressors were defined for the encoding period, based on the temporal structure (beat or interval) and modality (auditory or visual) of the conditions. Since each run consisted of the sequential presentation of the same task but different modalities (e.g. an auditory production task followed by a visual production task), the model thus included five regressors-of-interest for the three main tasks: (1) auditory beat, (2) auditory interval, (3) visual beat, (4) visual interval, and (5) response. In the NTFD random task runs, the encoding period additionally included an extra temporal structure—named random—which was also modelled separately for each modality. In this case, there were thus seven regressors-of-interest: (1) auditory beat, (2) auditory interval, (3) auditory random, (4) visual beat, (5) visual interval, (6) visual random, and (7) response. Depending on the task paradigm and the standard (for detailed information about the paradigms, refer to Methods and Figure 1), the duration of the unconvolved regressor’s plateau related to the encoding ranged from 3752 ms (for trials involving the shortest standard in the production task) to 6180 ms (for trials involving the longest standard and comparison in the perception task). The duration of the corresponding response regressor’s plateau varied based on the participant’s response time within a 2000 ms time window in all tasks. Rest was modelled implicitly.

To remove signal variance associated with low-frequency noise due to physiological factors or slow drifts in the fMRI signal, a set of discrete cosine transform functions was included as nuisance regressors in the design matrix for high-pass filtering, with a 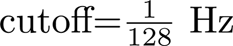.

Since our primary focus was on capturing the cognitive engagement involved in temporal encoding rather than evaluating participants’ success in the temporal estimation, regressors for bad performances were not taken into consideration. Moreover, the behavioural data indicated that the percentage of trials classified as bad or missing performances was negligible. In the production task, the mean and standard deviation of bad-trial percentages—calculated by first determining bad trials for each participant individually and then averaging these values across participants—was 4.49 *±* 7.22 %, comprising 3.10 *±* 6.94 % of missing trials and 1.38 *±* 0.80 % of trials where the mean-centred asynchronies exceeded three standard deviations. In the perception task, the mean and standard deviation of individual missing-trial percentages across all participants was 3.20 *±* 7.28 %, and no automatic-response trials (i.e. trials with a response time below 100 ms) were registered. Similarly, in the NTFD task, the mean and standard deviation of individual bad-trial percentages across participants was 8.36 *±* 8.54 %, comprising 2.18 *±* 5.68 % from missing trials, 2.80 *±* 3.56 % from automatic-response trials as well as responses exceeding 700 ms, and 3.75 *±* 2.49 % from incorrect discrimination, with a 0.36 *±* 0.57 % overlap between automatic-response and incorrect discrimination trials. In the NTFD random task, the mean and standard deviation of individual bad-trial percentages across participants was 7.53 *±* 7.52 %, comprising 0.97 *±* 2.67 % from missing trials, 3.74 *±* 5.21 % from automatic-response trials as well as responses exceeding 700 ms, and 3.17 *±* 2.49 % from incorrect discrimination, with a 0.35 *±* 0.80 % overlap between automatic-response and incorrect discrimination trials. For details about bad-trials estimation, consult Methods. We note that incorrect discrimination refers here to feature discrimination rather than the implicit temporal estimation involved in the task. Since the discrimination of the final event was straightforward (see Methods for details on the task paradigm), it served as an indicator of participants’ engagement, which was the relevant aspect to monitor in evaluating their performance.

#### Model Estimation and Individual Data Derivatives

For each subject, GLM-parameters estimation was restricted to voxels within functional brain regions by applying their individual whole-brain mask in native space—described in Methods—as an explicit mask, with an implicit masking threshold set to 1. To mitigate the effects of artefacts, we modelled noise using Robust Weighted Least Squares Estimation [59], which assigns lower weights to points in the time series exhibiting large residuals due to movement and physiological fluctuations. A mass-univariate GLM was fitted to the individual preprocessed functional data. Coefficient maps for regressors-of-interest (see Methods) were estimated and linearly contrasted to capture differences in evoked responses between a regressor-of-interest and rest or between two regressors-of-interest. For each subject, we obtained both single-run contrast estimates and multi-run fixed-effects contrast estimates, weighted by the number of runs across the two sessions. All main results are based on the fixed-effects estimates, whereas run-specific estimates were used only for the reliability analyses (see Methods). Corresponding PSC maps were additionally computed to correct for inflation of regression estimates associated with the shorter duration of the response period relative to encoding-related regressors. For the subsequent volume-based analyses, all individual unthresholded maps were normalized in the standard MNI152 space, and masked with a normalized whole-brain group mask. GLM estimation and generation of volume-based data derivatives were conducted using SPM12.

Surface-based analyses used the non-normalized, unthresholded individual contrast maps. For each participant, the volumetric contrast was sampled onto the fs LR32k surface by averaging along the cortical ribbon joining the individual white and pial surfaces (*vol to surf*, Nilearn v0.11.1 [60]); the midpoint of this ribbon defines the *midthickness*.

#### Summary Statistics

Group-level statistical maps were obtained by performing, for each contrast, a one-sample *t*-test on the individual contrast estimates at every voxel (or vertex), followed by a *t*-to-*Z* transform. Before this test, each participant’s contrast was smoothed with an 8 mm FWHM kernel: in MNI space within the group whole-brain mask for volume-based maps (whole-brain, dorsal striatum, and cerebellum), and geodesically along the fs LR32k midthickness surface for surface-based maps.

The cerebellum group map was then projected onto the *Spatially Unbiased Infratentorial Template* (SUIT) [61] using the SUITPy toolbox [62] v1.3.2, and the dorsal striatum group map onto a 26k-vertex surface mesh of this region (see Methods); neither projection involved additional smoothing.

For the group activation maps displayed in Figure 2, cortical results were computed on the fs LR32k surface to respect cortical folding, whereas cerebellar and striatal results were computed in volume; all peak-level (Extended Data Table E1) and subsequent ROI-based analyses were performed in volume for both cortical and subcortical regions.

*False Discovery Rate* (FDR) correction was applied using the *Benjamini–Hochberg* (BH) procedure, with BH-adjusted *p*-values computed for every voxel across the whole-brain volume, as implemented in *Nilearn*.

#### Dorsal-Striatum Volume-to-Surface Projection

We constructed the left and right rostrodorsalstriatum GIfTI meshes by combining coordinate and topology files obtained from the *Computerized Anatomical Reconstruction and Editing Tool Kit* (CARET, [63]). To improve mesh tessellation for mapping and visualization, vertex density was increased through triangle subdivision and subsequent Laplacian smoothing, reducing local surface irregularities while preserving global shape, using *trimesh* v4.6.6 (https://github.com/mikedh/trimesh). Afterwards, we generated an inner striatal surface by uniformly scaling the surface inward around its centroid: vertices were translated to the mesh centroid, scaled, and translated back, keeping topology fixed and ensuring that the inner surface remained concentric with the outer mesh.

This construction defines a striatal ribbon fully contained within the anatomical extent of the dorsal striatum, thereby avoiding sampling across boundaries with neighbouring structures. In addition, projection depth is constrained to a peri-striatal layer, allowing e.g. selective capture of putaminal signal within the lentiform nucleus.

Volume-to-surface mapping followed a line-sampling approach analogous to ribbon-constrained cortical projections [55], adapted here for the rostrodorsal striatum. For each surface vertex, a straight line was defined between the corresponding outer and inner striatal mesh vertices in MNI space, and a fixed number of points were generated uniformly along this line. These points define the sampling locations in continuous MNI space. Because sampling is restricted to line segments entirely contained within the striatal ribbon, the procedure is agnostic to signal originating from adjacent non-striatal regions. To extract volumetric activation values, each sampling location was transformed from MNI space into continuous voxel coordinates using the inverse affine of the NIfTI image. Voxel intensities were then estimated at these voxel-space locations via trilinear interpolation of the surrounding voxel grid, yielding one interpolated value per sample point. Finally, for each surface vertex, the maximum interpolated value across all sampled points along the line was retained, resulting in a single activation value assigned to that vertex. This analysis was performed using *SciPy* v1.15.2 [64].

Surface renderings were generated using *Plotly* 3D graphics library [65]. The refined striatal surface mesh was displayed with a greyscale background derived from surface curvature, serving as a geometric reference. Different surface projections were then overlaid onto this mesh.

#### Estimation of Individual ROI Masks

For the individualization of ROI masks within a given region, the process began with the creation of a group ROI mask. This was generated by intersecting the group-level encoding *vs.* rest contrast, thresholded at *p ≤* 0.001, with the corresponding anatomical atlas mask. This contrast was defined as the average of the auditory beat *vs.* rest, visual beat *vs.* rest, auditory interval *vs.* rest, and visual interval *vs.* rest, collapsed across all main tasks (production, perception, and NTFD). The resulting intersection defined the group ROI mask and its associated voxel count. Next, to derive individual ROI masks, the individual encoding map—also collapsed across all main tasks—was restricted to the corresponding atlas mask. Within this atlas-constrained map, the voxels with the highest activation values were selected such that the individualized ROI matched the voxel count of the group ROI mask. If the resulting individualized mask contained fewer voxels than the group ROI mask, a constrained binary non-isotropic dilation procedure was applied within the group ROI boundaries until the target voxel count was reached. Conversely, if more voxels than required were present after dilation, the lowest-valued voxels were iteratively removed to exactly match the group ROI size. In this way, the individualized ROIs were derived exclusively from individual encoding maps while still preserving the voxel count of the corresponding group ROI mask, and thereby ensuring identical ROI sizes across participants. The group-level and individual encoding maps used to define these ROI masks were spatially smoothed (8 mm FWHM), making voxel selection and peak localization more robust to noise. PSC values were subsequently extracted from unsmoothed maps (see Supplementary Note 2) to preserve ROI specificity; averaging across voxels within each ROI improved signal stability without introducing additional spatial smoothing [66, 67].

Because the location of functionally homologous regions varies across individuals in normalized space, group-or anatomically defined ROIs pool functionally heterogeneous voxels and blur condition-specific responses; defining ROIs within each participant attenuates this spatial mismatch and thereby increases both the sensitivity and the functional resolution of multi-subject analyses [22, 23]. To evaluate the influence of this individualization on our results, we repeated our key analyses using a purely group-based ROI for each region. In this case, the group ROI mask defined above—obtained by intersecting the thresholded group-level encoding *vs.* rest contrast with the corresponding atlas mask—was applied identically to all participants, with no subject-specific individualization. Group-based ROIs preserved the broad organization of our effects but reduced sensitivity to finer condition-specific differences, motivating the use of subject-specific ROIs throughout (see Methods).

Atlas masks for the auditory and visual cortices were both derived from the probabilistic Harvard-Oxford Cortical Structural Atlas [25, 26], using a probability threshold of 25%. Specifically, the auditory cortex mask was created by combining Heschl’s Gyrus, the *Planum Temporale* (PT), and the posterior division of the *Superior Temporal Gyrus* (STG). The visual cortex mask was similarly created by combining the Occipital Pole, the Occipital Fusiform Gyrus, and the superior and inferior subdivisions of the Lateral Occipital Cortex. In turn, the dorsal striatum mask was created by combining the Putamen and Caudate masks from the probabilistic Harvard-Oxford Subcortical Structural Atlas [29], using the same probability threshold. The cerebellum-atlas masks were created by combining masks of regions from the superior and inferior quadrants (tertiary and vermal areas were excluded) obtained from the Symmetric 128-region parcellation of the Nettekoven 2024 probabilistic atlas [28]. The atlas masks from premotor areas were directly obtained from the *Human Motor Area Template* (HMAT; [24]). In all cases, masks were defined bilaterally, i.e. encompassing the two hemispheres.

Anatomical labels for non-premotor cortical regions were derived from the Harvard–Oxford Cortical Structural Atlas. Functional labels for auditory and visual cortices were assigned with reference to the *Jülich Histological Atlas* (JHA) [68] and the *Probabilistic Maps of Visual Topography* (PMVT) [69], respectively. The cerebellum was functionally labelled using the Nettekoven 2024 hierarchical atlas described above. Striatal regions were labelled using both anatomical and connectivity-based criteria: anatomical labels came from the Harvard–Oxford Subcortical Structural Atlas, whereas functional labels followed the *Oxford-Imanova Striatal Connectivity Atlas* (OISCA) [30]. The striatal regions reported here corresponded to the *Executive* and *Rostral Motor* territories, reflecting dominant connectivity with regions of the prefrontal cortex associated with executive processing and motor planning/control, respectively (see Extended Data Table E1).

#### ROI Analyses

PSC for each region was extracted from unsmoothed individual contrast maps using individualized ROIs. These ROIs were functionally defined using the encoding *vs.* rest contrast, as defined in Methods. The extracted data represented the mean effect size, expressed as PSC, averaged across voxels within each ROI.

We then conducted a series of two-way RM ANOVAs across subjects on the encoding-period PSC relative to rest, each derived from the four regressors-of-interest defined as auditory beat, auditory interval, visual beat and visual interval (Methods). These analyses were either task-specific or pooled across tasks. For the NTFD-random conditions (Results), we isolated temporal predictability by defining a non-random contrast as the average of the beat and interval regressors (relative to rest) and comparing it with the random condition. PSC was extracted from the resulting auditory non-random, auditory random, visual non-random, and visual random contrasts. We also averaged the PSC across these four contrasts to obtain the mean response of each ROI, and tested it against zero with two-sided one-sample *t*-tests, Holm–Bonferroni-corrected across the eight ROIs. The encoding *vs.* rest contrast used to define the ROIs reflects the grand mean of the encoding period, collapsed across sensory modalities, tasks, and temporal structures. Because this grand mean is the sum of the four encoding regressors, it is orthogonal to the contrasts among them, so selecting voxels on it does not bias the condition effects tested in the main-task designs [70]. For the NTFD-random analyses, the same ROI definition was retained, keeping ROI delineation independent of the non-random and random effects, which were estimated from separate runs.

These analyses were performed using Nilearn v0.10.4, SciPy v1.13.1 [64], and pingouin v0.5.5 [71]. Omnibus *F*-tests were assessed using Greenhouse–Geisser correction, with the corresponding *ɛ* reported to indicate the magnitude of the sphericity correction: values close to 1 indicate little departure from sphericity, in which case corrected and uncorrected *p*-values converge. Significant omnibus effects were followed by post hoc pairwise *t*-tests, corrected using the Holm–Bonferroni step-down procedure to control the family-wise error rate at *α*_FWE_ = 0.05. Thus, for three pairwise task comparisons, the first step corresponded to *p ≤* 0.05*/*3, with subsequent steps evaluated against *p ≤* 0.05*/*2 and *p ≤* 0.05, respectively. For significant effects, we report uncorrected *p*-values that survived omnibus testing and, where applicable, Holm–Bonferroni correction. For visualization, central tendency was summarized using the median, with uncertainty estimated via percentile bootstrap confidence intervals based on 5000 subject-wise resamples.

#### Quantifying ROI Similarity with Between-Subject Correlation and Multidimensional Scaling

Similarity between ROIs was quantified by correlating ROI-specific activation vectors, each obtained by averaging PSC across the voxels of an ROI and concatenating the resulting condition-wise estimates. These vectors comprised the 12 main-task conditions (production, perception, and NTFD × beat and interval × auditory and visual) together with a synthetic rest condition (PSC = 0), yielding 13 activation profiles per ROI and subject; the synthetic rest condition provided a common baseline across ROIs. For each subject, similarity was computed as the *Pearson* correlation between the activation profiles of each ROI pair (*r_s_*). These coefficients were Fisher *Z*-transformed for group-level inference, whereas the raw *r_s_* values were retained to construct the ROI × ROI similarity matrix used for visualization and the MDS described below. Group-level inference used one-sample *t*-tests against zero for each of the 28 ROI pairs, Holm–Bonferroni corrected to control the family-wise error rate (*α*_FWE_ = 0.05, where, within the 28 pairwise tests, the smallest *p*-value was evaluated against *p ≤* 0.05*/*28). In addition, paired *t*-tests compared each premotor region’s subject-wise correlation with the cerebellum against that with the dorsal striatum, also Holm–Bonferroni corrected (*α*_FWE_ = 0.05, where, within the four tests, the smallest *p*-value was evaluated against *p ≤* 0.05*/*4).

Next, the resulting ROI × ROI similarity matrix was explored using classical *Multidimensional Scaling* (MDS). The correlation matrix was treated as a similarity matrix and normalized by its matrix rank prior to decomposition. Classical MDS was then applied to obtain a low-dimensional geometric representation of ROIs, in which distances between points reflect their pairwise similarity structure, with smaller distances indicating more similar activation profiles and larger distances indicating greater dissimilarity. The decomposition was performed using the standard eigendecomposition-based solution for classical MDS, as implemented in the *Pattern Component Modeling* toolbox [72]. Specifically, a three-component decomposition was examined to visualize the dominant dimensions of the ROI similarity structure in low-dimensional space.

#### Reliability Analysis

To quantify the stability of condition-specific activation profiles, we estimated withinsubject reliability using a split-half approach applied to run-specific GLM estimates. Condition-wise activation profiles were constructed as described in Methods. For each participant, PSC values were extracted from individualized ROIs (see Methods) and organized into vectors across experimental conditions, providing independent observations across runs. Reliability was estimated by partitioning the data into independent halves based on runs and computing, for each subject and ROI, the *Pearson* correlation between condition-wise activation profiles derived from the two halves. This procedure was repeated across all possible split-half combinations, yielding subject-level reliability estimates for each ROI. Split-half correlations were converted to full-length reliability estimates using the *Spearman–Brown prediction formula*. The approach described herein provides an empirical estimate of ROI reliability that informs the interpretation of differences in correlation strength between regions, relative to their measurement precision [73].

## Acknowledgements

We thank Scott Charlton for his assistance at the scanner throughout a demanding acquisition schedule, Suzanne Witt for curating the imaging data into the BIDS format, Aaron Gibbings for assisting with protocol validation, and Haitao Yang for providing technical support with hardware configuration and server access. We thank our colleagues at the Western Centre for Brain and Mind for their insightful feedback, particularly Ladan Shahshahani, Joshua Hoddinott, Marco Emanuele, and Ehsan Tousi. We also thank Verena Rafael, Katsiaryna Buchko, Silvana Medina, and Eduardo Allard for their assistance with behavioural-data collection.

## Author Contributions

A.L.P., J.D. and J.A.G. conceived the study and designed the task paradigms. A.L.P. developed the task protocols; A.L.P. and A.A. carried out their implementation. A.L.P. and V.K-A. collected the imaging data. A.L.P., V.K-A., A.A. and N.P. collected the behavioural data. A.L.P. curated and analysed the imaging and behavioural data; V.K-A. contributed to the preprocessing of the imaging data. J.D. and J.A.G. advised on data collection and analysis. A.L.P., J.D. and J.A.G. wrote the manuscript. All authors reviewed and approved the final version.

## Funding

This research was supported by the BrainsCAN initiative at Western University, funded by the Canada First Research Excellence Fund (CFREF), including a Postdoctoral Fellowship to A.L.P.; by an NSERC Discovery Grant (RGPIN/05869-2024NSERC) and an EWR Steacie Award (566202-2021-SMFSU) to J.A.G.; and by a CIHR Project Grant (PJT-191815) to J.D.

## Competing Interests

The authors declare no competing interests.

## Additional Information

### Ethics Approval and Consent to Participate

All participant-related procedures reported in this study were approved by the Health Sciences Research Ethics Board at Western University (Protocol #107293 for imaging sessions and Protocol #106385 for behavioural training sessions). Informed written consents were also obtained from each participant before the first behavioural and imaging sessions, in accordance with the Declaration of Helsinki.

### Code Availability

Documented metadata, which concern the programmed sequence of stimuli presented during behavioural and imaging sessions, were made publicly available on https://github.com/alpinho/tdtb_protocols. They include: (1) the behavioural, imaging, and practice versions of the protocols for all tasks; (2) instructions to the participants; (3) generation and randomization of the stimuli; and (4) demo presentations of the tasks as video files.

The scripts used for both behavioural and imaging data analysis were also made available through GitHub under the BSD 3-Clause License: https://github.com/alpinho/tdtb analysis.

## Extended Data

**Figure E1.**
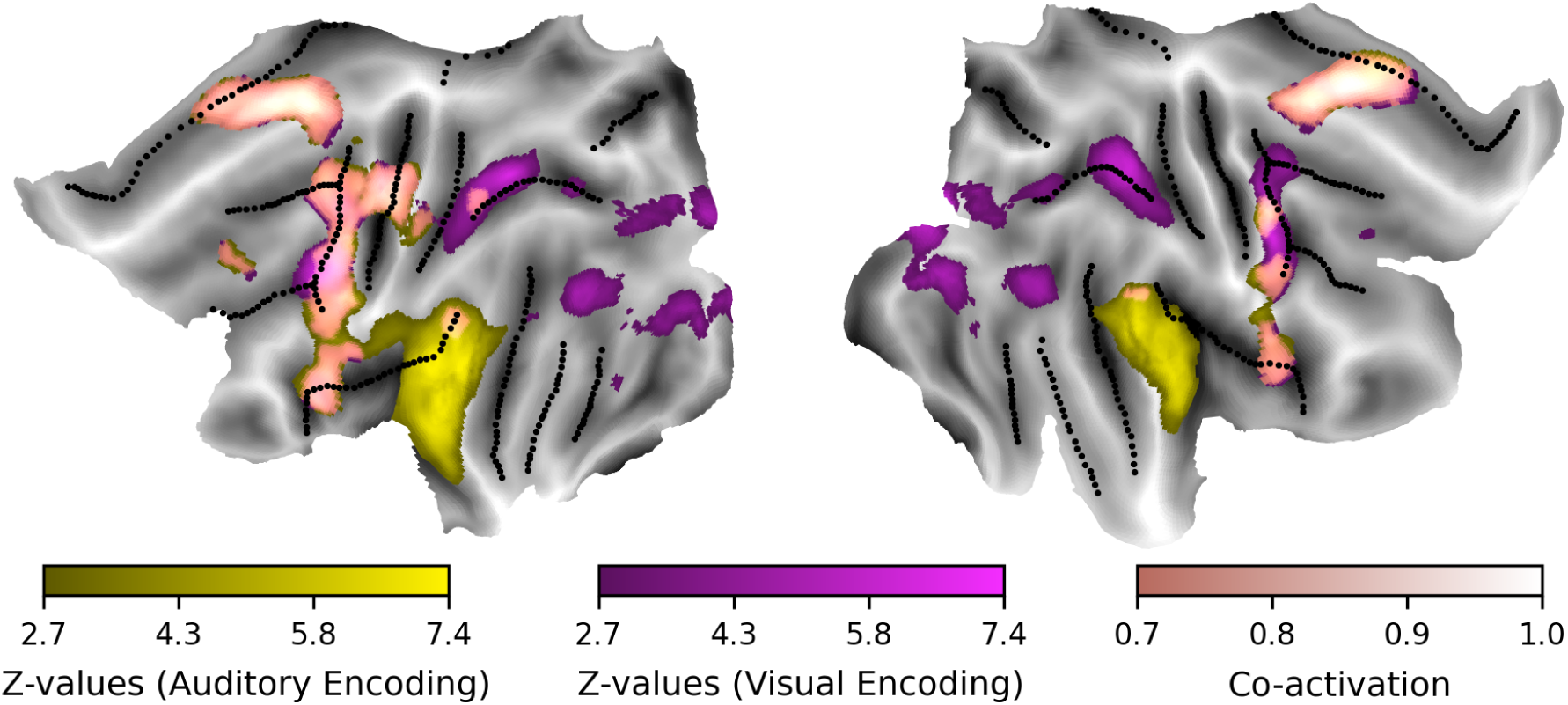
Modality-specific sensory recruitment during temporal encoding. Group-level *Z*-maps of the encoding period computed separately for auditory and visual tasks, thresholded using a whole-brain FDR correction (*q* = 0.05). The auditory encoding *vs.* rest contrast shows bilateral activation centred on Heschl’s gyrus, including the *Primary Auditory Cortex* (TE1.0, TE1.1 and TE1.2) as defined by the JHA [68], with extensions towards PT and the posterior division of the STG. The visual encoding *vs.* rest contrast reveals activation in the Occipital Pole and the Occipital Fusiform Gyrus, spanning dorsal and ventral territories of the *Primary Visual Cortex* (V1–V3) and the *Early Ventral Visual Cortex* (hV4), defined according to the Probabilistic Maps of Visual Topography [69], as well as the Superior Parietal Lobe. Vertices concurrently active in both contrasts are shown in the co-activation map, defined as the voxel-wise sum of the two normalized *Z*-maps. The colour bar spans 0.7 to 1.0, where the lower bound corresponds to both maps at their FDR threshold and the upper bound to at least one map at its peak. Sensory-specific recruitment is thus confined to the primary and sensory cortices, whereas the co-activation extends beyond them and is centred on premotor regions, preserving the broader cortico–striatal–cerebellar architecture of temporal encoding.

**Figure E2.**
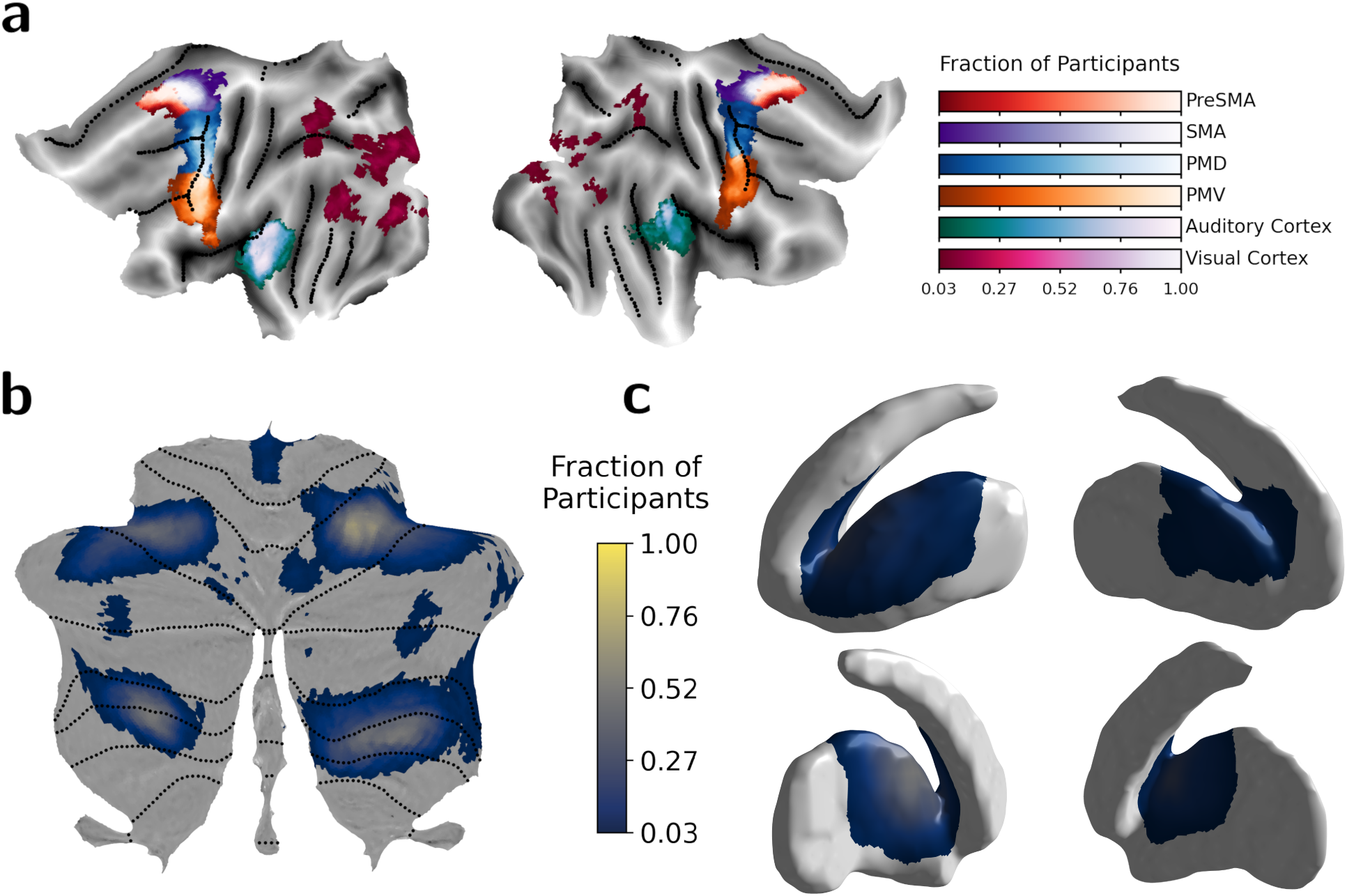
Individualized non-contiguous ROIs defined as a fraction of participants represented at every vertex. Maps represent eight volume-based ROIs, each defined bilaterally in MNI space and individualized for every participant by combining that participant’s encoding *vs.* rest contrast with atlas-based regions. This follows frameworks that define ROIs functionally within each subject while constraining them with group-level information [22, 75], extended here into a functional– anatomical strategy. Values at each vertex denote the fraction of participants whose individualized ROI includes that location. **(a)** flat projection onto fs LR32k of the four premotor and two sensory cortical ROIs (PMD, PMV, SMA and PreSMA; auditory and visual cortices); **(b)** flat projection of the cerebellar ROI onto SUIT space, drawn from the *Demand* functional domain [28]; and **(c)** projection of the striatal ROI onto the rostrodorsal-striatum surface, comprising anterior and mainly medial portions of the Putamen with extensions towards the head of the Caudate Nucleus. Sections **b** and **c** share the same colour bar. The derivation procedure is described in Methods.

**Figure E3.**
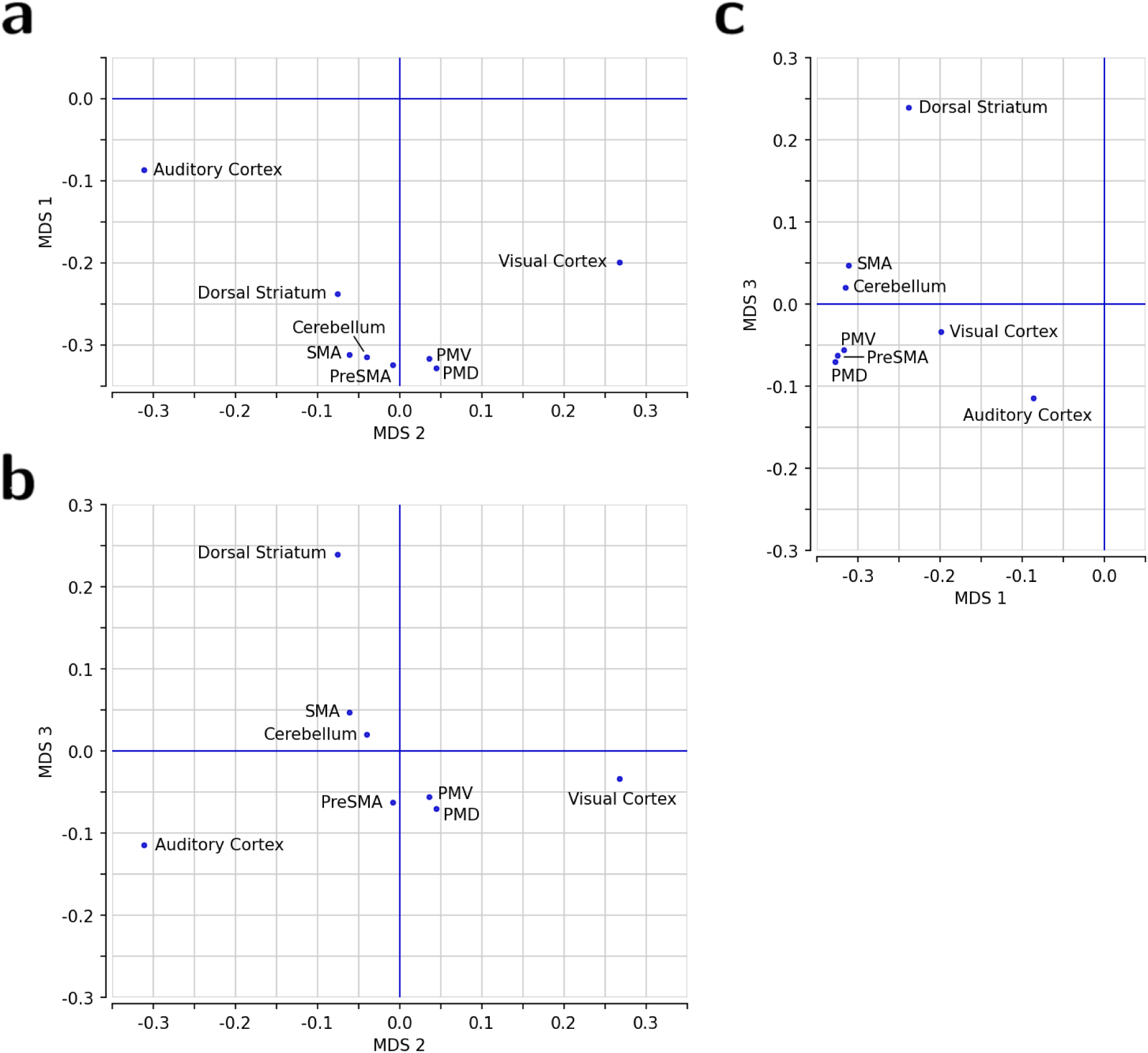
Two-dimensional projections of the ROI similarity structure obtained with classical MDS. Projections are derived from a three-dimensional MDS decomposition applied to the between-subject matrix of correlations between the encoding-period PSCs of ROI pairs, shown in three dimensions in Figure 3b. Each point represents an ROI, and distances between points depict their functional similarity across experimental conditions. Panels show the three pairwise combinations of the MDS dimensions: **(a)** MDS 2 *vs.* MDS 1; **(b)** MDS 2 *vs.* MDS 3; and **(c)** MDS 1 *vs.* MDS 3. Axes are ordered by decreasing eigenvalue magnitude. MDS 1 captures variability shared across all regions, grouping the premotor regions (PreSMA, SMA, PMD and PMV) with the cerebellum; MDS 2 reflects variability associated primarily with the sensory cortices, separating the auditory and visual cortices from that premotor–cerebellar cluster; and MDS 3 isolates the dorsal striatum, reflecting task-related variability that distinguishes it from the cortical and cerebellar regions.

**Table E1.**
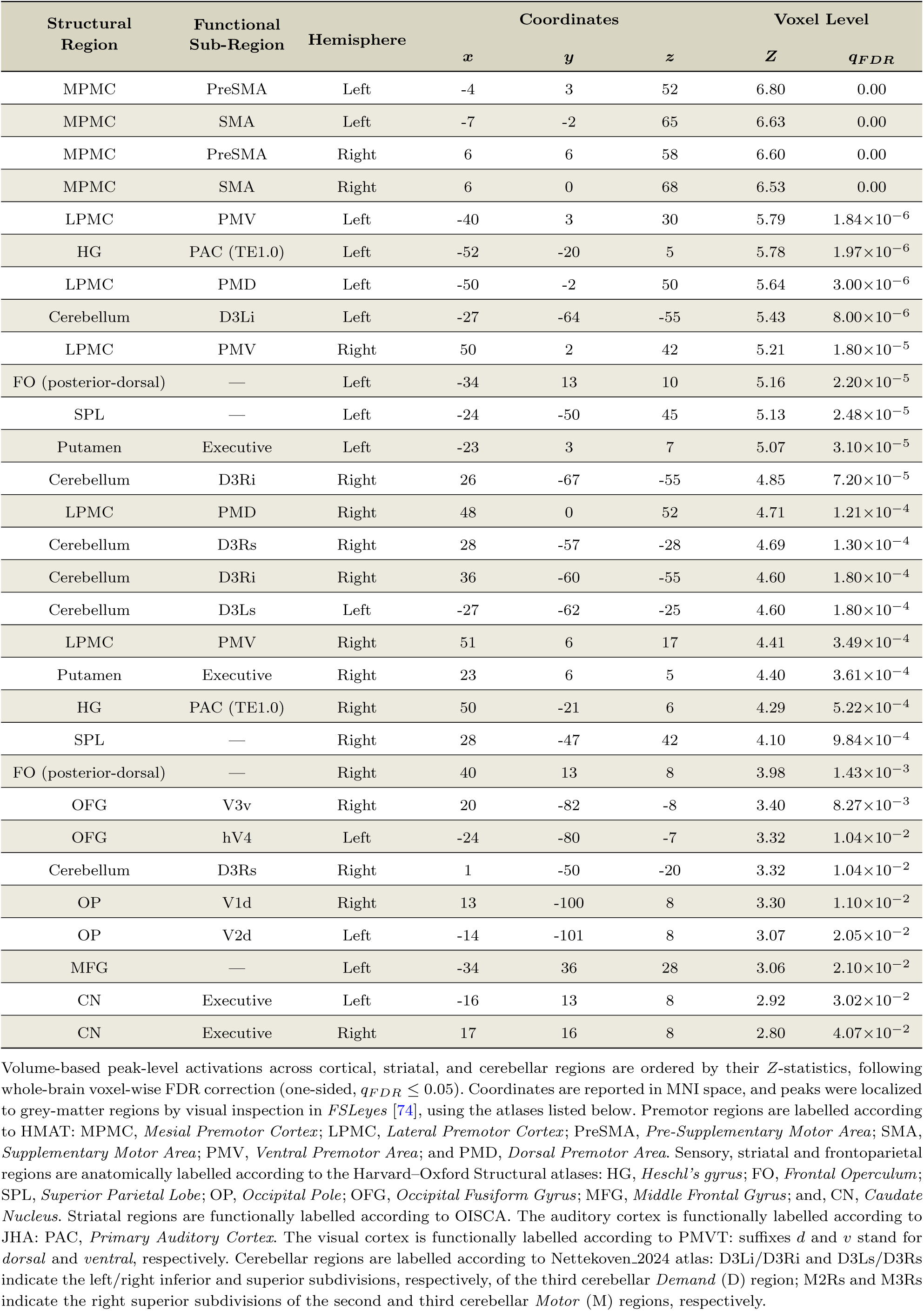
Imaging results for the contrast encoding *vs.* rest.

| Structural Region | Functional Sub-Region | Hemisphere | Coordinates |  |  | Voxel Level |  |
|---|---|---|---|---|---|---|---|
|  |  |  | *x* | *y* | *z* | *Z* | *q*<sub>*FDR*</sub> |
| MPMC | PreSMA | Left | -4 | 3 | 52 | 6.80 | 0.00 |
| MPMC | SMA | Left | -7 | -2 | 65 | 6.63 | 0.00 |
| MPMC | PreSMA | Right | 6 | 6 | 58 | 6.60 | 0.00 |
| MPMC | SMA | Right | 6 | 0 | 68 | 6.53 | 0.00 |
| LPMC | PMV | Left | -40 | 3 | 30 | 5.79 | 1.84×10<sup>−6</sup> |
| HG | PAC (TE1.0) | Left | -52 | -20 | 5 | 5.78 | 1.97×10<sup>−6</sup> |
| LPMC | PMD | Left | -50 | -2 | 50 | 5.64 | 3.00×10<sup>−6</sup> |
| Cerebellum | D3Li | Left | -27 | -64 | -55 | 5.43 | 8.00×10<sup>−6</sup> |
| LPMC | PMV | Right | 50 | 2 | 42 | 5.21 | 1.80×10<sup>−5</sup> |
| FO (posterior-dorsal) | — | Left | -34 | 13 | 10 | 5.16 | 2.20×10<sup>−5</sup> |
| SPL | — | Left | -24 | -50 | 45 | 5.13 | 2.48×10<sup>−5</sup> |
| Putamen | Executive | Left | -23 | 3 | 7 | 5.07 | 3.10×10<sup>−5</sup> |
| Cerebellum | D3Ri | Right | 26 | -67 | -55 | 4.85 | 7.20×10<sup>−5</sup> |
| LPMC | PMD | Right | 48 | 0 | 52 | 4.71 | 1.21×10<sup>−4</sup> |
| Cerebellum | D3Rs | Right | 28 | -57 | -28 | 4.69 | 1.30×10<sup>−4</sup> |
| Cerebellum | D3Ri | Right | 36 | -60 | -55 | 4.60 | 1.80×10<sup>−4</sup> |
| Cerebellum | D3Ls | Left | -27 | -62 | -25 | 4.60 | 1.80×10<sup>−4</sup> |
| LPMC | PMV | Right | 51 | 6 | 17 | 4.41 | 3.49×10<sup>−4</sup> |
| Putamen | Executive | Right | 23 | 6 | 5 | 4.40 | 3.61×10<sup>−4</sup> |
| HG | PAC (TE1.0) | Right | 50 | -21 | 6 | 4.29 | 5.22×10<sup>−4</sup> |
| SPL | — | Right | 28 | -47 | 42 | 4.10 | 9.84×10<sup>−4</sup> |
| FO (posterior-dorsal) | — | Right | 40 | 13 | 8 | 3.98 | 1.43×10<sup>−3</sup> |
| OFG | V3v | Right | 20 | -82 | -8 | 3.40 | 8.27×10<sup>−3</sup> |
| OFG | hV4 | Left | -24 | -80 | -7 | 3.32 | 1.04×10<sup>−2</sup> |
| Cerebellum | D3Rs | Right | 1 | -50 | -20 | 3.32 | 1.04×10<sup>−2</sup> |
| OP | V1d | Right | 13 | -100 | 8 | 3.30 | 1.10×10<sup>−2</sup> |
| OP | V2d | Left | -14 | -101 | 8 | 3.07 | 2.05×10<sup>−2</sup> |
| MFG | — | Left | -34 | 36 | 28 | 3.06 | 2.10×10<sup>−2</sup> |
| CN | Executive | Left | -16 | 13 | 8 | 2.92 | 3.02×10<sup>−2</sup> |
| CN | Executive | Right | 17 | 16 | 8 | 2.80 | 4.07×10<sup>−2</sup> |

Volume-based peak-level activations across cortical, striatal, and cerebellar regions are ordered by their *Z*-statistics, following whole-brain voxel-wise FDR correction (one-sided, *q*<sub>*FDR*</sub> ≤ 0.05). Coordinates are reported in MNI space, and peaks were localized to grey-matter regions by visual inspection in *FSLeyes* [74], using the atlases listed below. Premotor regions are labelled according to HMAT: MPMC, *Mesial Premotor Cortex*; LPMC, *Lateral Premotor Cortex*; PreSMA, *Pre-Supplementary Motor Area*; SMA, *Supplementary Motor Area*; PMV, *Ventral Premotor Area*; and PMD, *Dorsal Premotor Area*. Sensory, striatal and frontoparietal regions are anatomically labelled according to the Harvard–Oxford Structural atlases: HG, *Heschl’s gyrus*; FO, *Frontal Operculum*; SPL, *Superior Parietal Lobe*; OP, *Occipital Pole*; OFG, *Occipital Fusiform Gyrus*; MFG, *Middle Frontal Gyrus*; and, CN, *Caudate Nucleus*. Striatal regions are functionally labelled according to OISCA. The auditory cortex is functionally labelled according to JHA: PAC, *Primary Auditory Cortex*. The visual cortex is functionally labelled according to PMVT: suffixes *d* and *v* stand for *dorsal* and *ventral*, respectively. Cerebellar regions are labelled according to Nettekoven\_2024 atlas: D3Li/D3Ri and D3Ls/D3Rs indicate the left/right inferior and superior subdivisions, respectively, of the third cerebellar *Demand* (D) region; M2Rs and M3Rs indicate the right superior subdivisions of the second and third cerebellar *Motor* (M) regions, respectively.

**Table E2.**
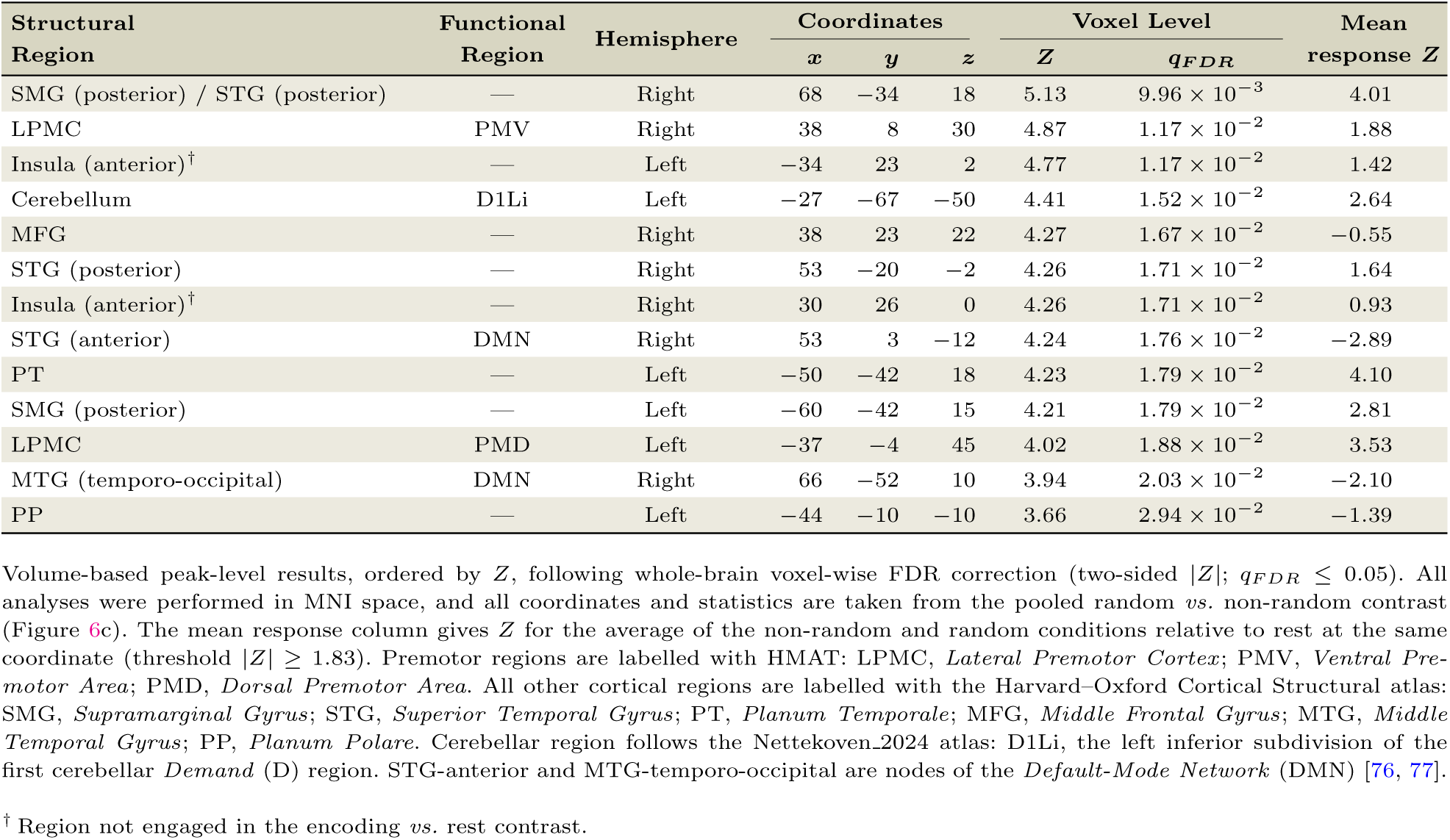
Imaging results for the contrast random *vs.* non-random.

| Structural Region | Functional Region | Hemisphere | Coordinates |  |  | Voxel Level |  | Mean response *Z* |
|---|---|---|---|---|---|---|---|---|
|  |  |  | *x* | *y* | *z* | *Z* | *q*<sub>*FDR*</sub> |  |
| SMG (posterior) / STG (posterior) | — | Right | 68 | −34 | 18 | 5.13 | 9.96 × 10<sup>−3</sup> | 4.01 |
| LPMC | PMV | Right | 38 | 8 | 30 | 4.87 | 1.17 × 10<sup>−2</sup> | 1.88 |
| Insula (anterior)<sup>†</sup> | — | Left | −34 | 23 | 2 | 4.77 | 1.17 × 10<sup>−2</sup> | 1.42 |
| Cerebellum | D1Li | Left | −27 | −67 | −50 | 4.41 | 1.52 × 10<sup>−2</sup> | 2.64 |
| MFG | — | Right | 38 | 23 | 22 | 4.27 | 1.67 × 10<sup>−2</sup> | −0.55 |
| STG (posterior) | — | Right | 53 | −20 | −2 | 4.26 | 1.71 × 10<sup>−2</sup> | 1.64 |
| Insula (anterior)<sup>†</sup> | — | Right | 30 | 26 | 0 | 4.26 | 1.71 × 10<sup>−2</sup> | 0.93 |
| STG (anterior) | DMN | Right | 53 | 3 | −12 | 4.24 | 1.76 × 10<sup>−2</sup> | −2.89 |
| PT | — | Left | −50 | −42 | 18 | 4.23 | 1.79 × 10<sup>−2</sup> | 4.10 |
| SMG (posterior) | — | Left | −60 | −42 | 15 | 4.21 | 1.79 × 10<sup>−2</sup> | 2.81 |
| LPMC | PMD | Left | −37 | −4 | 45 | 4.02 | 1.88 × 10<sup>−2</sup> | 3.53 |
| MTG (temporo-occipital) | DMN | Right | 66 | −52 | 10 | 3.94 | 2.03 × 10<sup>−2</sup> | −2.10 |
| PP | — | Left | −44 | −10 | −10 | 3.66 | 2.94 × 10<sup>−2</sup> | −1.39 |

Volume-based peak-level results, ordered by *Z*, following whole-brain voxel-wise FDR correction (two-sided |*Z*|; *q*<sub>*FDR*</sub> ≤ 0.05). All analyses were performed in MNI space, and all coordinates and statistics are taken from the pooled random *vs.* non-random contrast (Figure 6c). The mean response column gives *Z* for the average of the non-random and random conditions relative to rest at the same coordinate (threshold |*Z*| ≥ 1.83). Premotor regions are labelled with HMAT: LPMC, *Lateral Premotor Cortex*; PMV, *Ventral Pre-motor Area*; PMD, *Dorsal Premotor Area*. All other cortical regions are labelled with the Harvard–Oxford Cortical Structural atlas: SMG, *Supramarginal Gyrus*; STG, *Superior Temporal Gyrus*; PT, *Planum Temporale*; MFG, *Middle Frontal Gyrus*; MTG, *Middle Temporal Gyrus*; PP, *Planum Polare*. Cerebellar region follows the Nettekoven\_2024 atlas: D1Li, the left inferior subdivision of the first cerebellar *Demand* (D) region. STG-anterior and MTG-temporo-occipital are nodes of the *Default-Mode Network* (DMN) [76, 77].
<sup>†</sup> Region not engaged in the encoding *vs.* rest contrast.

## Supplementary Information

### Supplementary Note 1: Whole-Brain Effects of Temporal Regularity

Among the peaks where random sequences raised activity above rest, only the anterior insula lay outside the regions engaged during encoding; the others fell within the encoding *vs.* rest map or, where a peak lay just beyond it, in a region tabulated there (Extended Data Table E1). At the anterior insula the encoding response fell well below threshold (left *Z* = 1.71, right *Z* = 0.29, against *Z ≥* 2.72, one-sided as in Extended Data Table E1). There, non-random sequences left activity at baseline (left *Z* = −0.17, right *Z* = −0.86, against *|Z| ≥* 1.83), whereas random sequences raised it above rest (left *Z* = 2.78, right *Z* = 2.64, against *|Z| ≥* 1.89). The mean of the two conditions also stayed below threshold (left *Z* = 1.42, right *Z* = 0.93, against *|Z| ≥* 1.83), so the region appears in the difference map but not in the mean. The nearest encodingrelated site, the posterior-dorsal Frontal Operculum, lies 13 mm away on the left and 18 mm on the right. It was engaged during encoding (left *Z* = 5.16, right *Z* = 3.98), responded to both non-random (left *Z* = 4.44, right *Z* = 2.45) and random sequences (left *Z* = 4.92, right *Z* = 3.28), and did not distinguish them, though narrowly on the left (left *Z* = 2.920, right *Z* = 2.76, against *|Z| ≥* 2.923).

### Supplementary Note 2: Robustness of Results to ROI Definition

To assess the impact of ROI definition on the reported effects, we repeated the key analyses using only groupdefined ROIs instead of the subject-specific ones (see Methods). Subject-specific ROIs were the a priori choice, motivated by prior work showing that within-subject functional localization increases sensitivity and functional resolution [22, 23, 78]. This analysis therefore tests whether our conclusions are robust to ROI definition.

We first recomputed the inter-regional between-subject similarity matrix (Supplementary Fig. S1). Groupdefined ROIs preserved the overall organization obtained with individualized ROIs (Figure 3a), including the core cluster of high correlations among the premotor regions and the cerebellum. Almost all inter-regional correlations were weaker under the group definition. The reduction was largest for pairs involving the visual cortex, most notably PMD–visual cortex (*r_s_* from 0.62 to 0.33), and, within the core cluster, was concentrated on pairs involving the lateral premotor regions PMD and PMV (e.g. PMV–cerebellum, from 0.74 to 0.52, and PMD–PMV, from 0.85 to 0.66), whereas correlations among the medial premotor regions and their links to the cerebellum changed little (e.g. PreSMA–SMA, from 0.74 to 0.73). This reduction also brought four associations below significance under the group definition, i.e. dorsal striatum–visual cortex, SMA–visual cortex, and cerebellum–visual cortex, together with PMV–auditory cortex, which fell close to zero, whereas every pair remained significant with individualized ROIs. Correlations involving the auditory cortex were otherwise comparatively stable, the cerebellum–auditory cortex pair being the only one to increase (from 0.28 to 0.33). The single negative association, auditory cortex–visual cortex, was marginally stronger under the group definition.

We also tested whether the single dissociation between the dorsal striatum and the cerebellum during auditory beat-based encoding (Figure 5) depended on ROI definition. With group-defined ROIs, the ROI × Temporal Structure interaction for the auditory modality remained significant but was markedly attenuated relative to the individualized ROIs (*F*_(1,30)_ = 6.68, *p* = 0.015, *ɛ* = 1). Critically, the post hoc effect underlying the dissociation, i.e. higher dorsal-striatal activation for auditory beat than auditory interval, no longer survived correction (*t*_(30)_ = 2.06, *p* = 0.048, *d* = 0.10; Holm–Bonferroni corrected, *α*_FWE_ = 0.05, where, within the two ROI-specific tests, the smallest *p*-value was evaluated against *p ≤* 0.05*/*2), and the cerebellum again remained non-significant. Thus, although a weaker ROI × Temporal Structure interaction persists, the dorsal-striatal beat sensitivity that defines the single dissociation is reliably resolved only with subject-specific ROIs. The same held for the task-resolved effect in NTFD, which with group-defined ROIs lost both the interaction (*F*_(1,30)_ = 1.94, *p* = 0.174, *ɛ* = 1) and the dorsal-striatal post hoc (*t*_(30)_ = 1.80, *p* = 0.083, *d* = 0.15).

**Figure S1.**
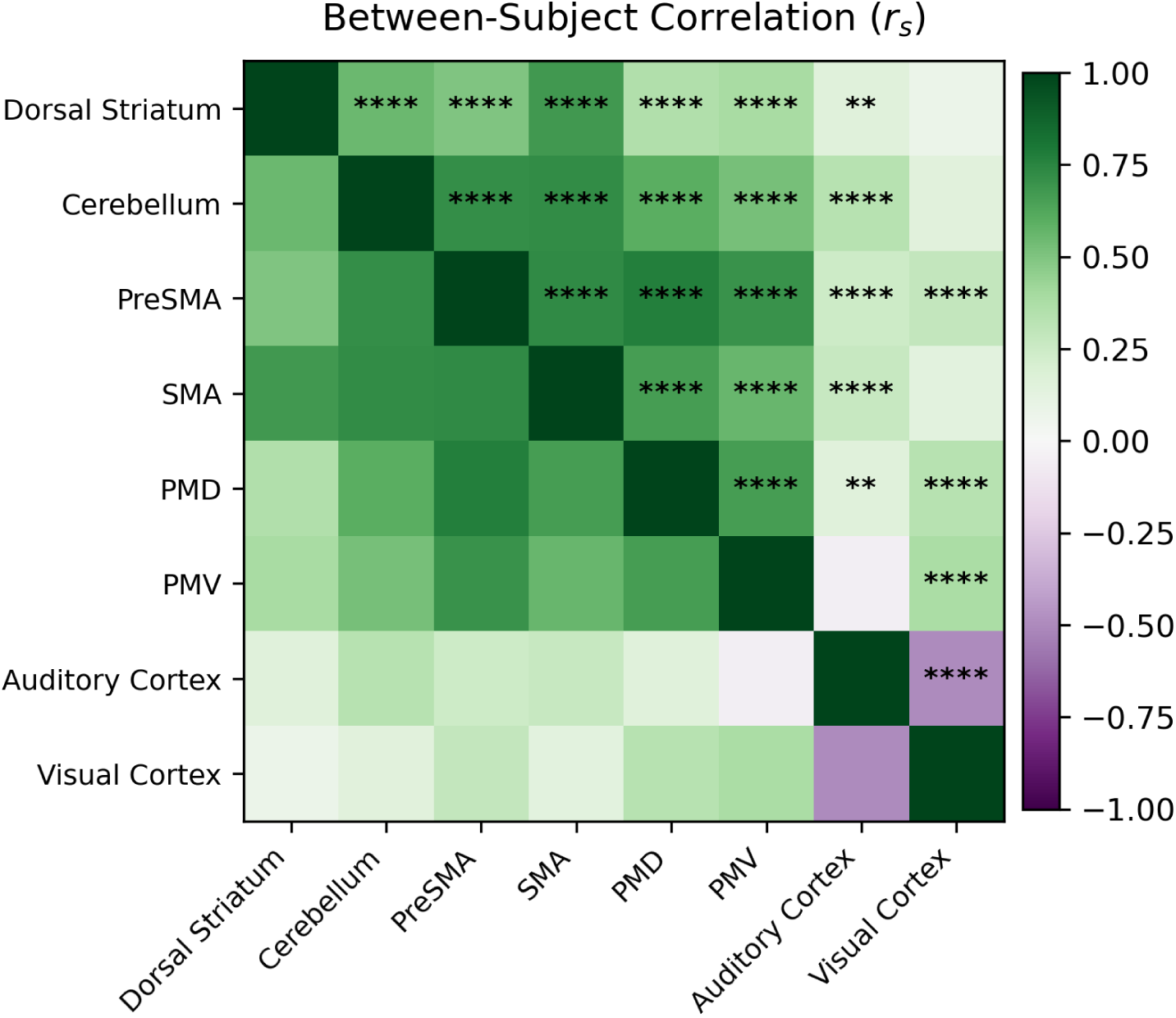
ROI-similarity structure obtained with group-defined ROIs. ROI-by-ROI similarity matrix computed from subject-wise correlations of condition-evoked PSC activation profiles. Each cell reports the mean *Pearson* correlation across subjects (*rs*), quantifying the extent to which activation profiles across experimental conditions covary between pairs of group-defined ROIs. Stars denote the uncorrected *p*-value at which the Fisher-*Z*-transformed correlations differ significantly from zero across subjects, and are shown only for ROI pairs surviving Holm–Bonferroni correction across the 28 pairs (*α*_FWE_ = 0.05; * *p <* 0.05, ** *p <* 0.01, *** *p <* 0.001, **** *p <* 10^−4^).

Overall, the coarse architecture replicates under either ROI definition, so the main organization does not depend on the ROI choice. The two definitions differ, however, in the finer, condition-specific structure. Groupdefined ROIs blur it: the inter-regional similarities weaken and the dorsal-striatal sensitivity to auditory beat is lost. This loss under group ROIs is the pattern expected for a less sensitive definition when functional regions are variably positioned across individuals [22, 23, 78].

### Supplementary Note 3: Trial-level Timing Fidelity of the Beat Condition

Methods summarizes the rendering deviations by their RMS, a measure of their typical size. Here, for the beat condition in both modalities, we report what the RMS cannot show: the fraction of trials on which a rendered duration exceeded a published threshold.

We used two bounds. The first is the duration-discrimination threshold, expressed as a *Weber Fraction*: the change in an empty duration that a listener detects reliably is about 5% of that duration [39]. The second is the pulse-attribution limit, i.e. the timing deviation, about 8.6%, beyond which a regular pulse can no longer be attributed to a sequence [40].

Both bounds are fractions of a duration, so we applied each to the element’s own nominal: the standards to *S*, the intervening SOAs to 3*S*. The choice matters for the intervening SOAs. Because the “beat” sequences are non-isochronous, an intervening SOA is a single sounded duration of 3*S* that spans pulse positions carrying no onset. Scaling its error by the pulse period *S* would measure it against those silent positions, as if the SOA were three-pulse durations; scaling by 3*S* measures it against the two onsets that actually bound it. Concretely, the standards give 23–33 ms at 5% and 39–57 ms at 8.6%; in turn, the SOAs give 69–99 ms at 5% and 118–171 ms at 8.6%.

The two bounds presuppose different things. The 5% bound concerns the durations alone and holds whether or not a beat is heard, so it measures duration fidelity. The 8.6% bound concerns a pulse, so it applies only if a beat is perceived. For the beat condition, that is likely but not certain: its integer-ratio structure favours a metrical percept [16, 17], while its sparsity weakens the pulse a listener would infer [16]. That same sparsity makes the bound approximate: sensitivity to a timing deviation depends on how many intervals a sequence provides [79], but Madison and Merker measured 8.6% on a denser, fully sounded sequence [40]. We therefore treated 8.6% as an approximate, beat-relevant reference: the fraction of trials on which a perceived pulse could be lost.

For a trial, let *s*_1_*, s*_2_ be the two rendered standards and *g*_1_*, g*_2_ the two rendered SOAs, with nominal durations *S* and 3*S* respectively, and let *f* be the fraction (0.05 or 0.086). Two criteria index accuracy, comparing each element to its nominal; two index internal consistency, comparing the paired elements to each other; and one indexes the ratio between the intervening SOAs and the standards, nominally 3. A trial is flagged when a criterion is met:

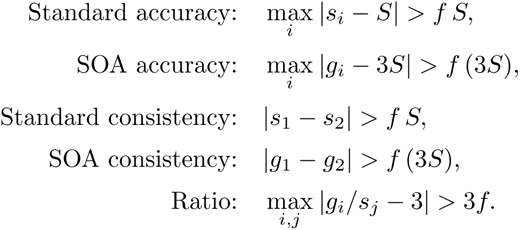

Each accuracy criterion uses the larger of the two elements’ deviations from nominal, whereas each consistency criterion uses the difference between the paired elements. The ratio criterion uses the largest deviation among all pairs of an intervening SOA and a standard of the trial. NTFD trials contain a third standard; for these, standard accuracy uses the largest deviation among the three standards, and standard consistency the range across them. Supplementary Table S1 reports the proportion of trials flagged by each criterion under both bounds.

**Table S1.** Percentage of beat trials with a timing deviation above each threshold.

|  | **Auditory** |  | **Visual** |  |
|---|---|---|---|---|
| **Regularity Check** | **> 5%** | **> 8.6%** | **> 5%** | **> 8.6%** |
| **Standard accuracy:** a standard deviates from its nominal *S* | 27.6% | 4.8% | 0.0% | 0.0% |
| **SOA accuracy:** an SOA deviates from its nominal 3*S* | 0.0% | 0.0% | 0.0% | 0.0% |
| **Standard consistency:** the standards disagree | 24.7% | 4.9% | 0.0% | 0.0% |
| **SOA consistency:** the two SOAs disagree | 0.0% | 0.0% | 0.0% | 0.0% |
| **Ratio:** an SOA-to-standard ratio deviates from 3 | 24.5% | 4.8% | 0.0% | 0.0% |

Thresholds are fractions of each element's own nominal (*S* for the standards, 3*S* for the intervening SOAs), or of the nominal ratio 3. The 5% columns are the duration-discrimination Weber Fraction [39]: 23–33 ms for standards, and 69–99 ms for SOAs. The 8.6% columns are the pulse-attribution limit [40]: 39–57 ms for standards, and 118–171 ms for SOAs. *N* = 330 auditory and 315 visual recordings (324 and 314 for standard consistency, which requires at least two valid standards).

In the auditory beat condition, no intervening SOA deviated from 3*S* by more than 5%, and the two SOAs never differed by more than 5%. Against the 5% bound, about a quarter of trials contained a standard that deviated from its nominal, and the standards differed from each other about as often; against the 8.6% bound, about 5% of trials contained a deviation this large. The ratio between the intervening SOAs and the standards, which defines the beat’s integer-ratio structure, deviated from 3 about as often (24.5% and 4.8% of trials), and only on trials in which a standard was also flagged. A perceived beat, if one is induced, could therefore have been compromised on these trials, which cannot be identified, and possibly beyond them (Methods). In the visual beat condition, no trial exceeded either bound on any criterion. Because the nominal timing matches the intended metrical structure exactly, these deviations can only reduce the beat’s regularity, never raise it. Thus, they can only weaken a regularity-driven effect, not give rise to one, as argued in Methods.

### Supplementary Note 4: Consistency of Behavioural Performance across Cohorts

The auditory protocols were re-implemented in PsychoPy and used to collect a second cohort, which renders the auditory timing with substantially less jitter than Expyriment (Methods). To test whether this change of presentation software altered behaviour, we compared performance between the two cohorts on the three tasks, using the auditory data from the first two behavioural sessions. Each task contributed one metric per participant and temporal structure (Methods): the Mean SA for the production task, the DL for the perception task, and the RT for the NTFD task. All 31 first-cohort participants, who are those with imaging data (see Methods), completed both sessions; in the second cohort, 17 completed one session and 16 of them completed two, and each metric was aggregated per participant over the completed sessions.

The comparison is a 2 × 2 design with one between-subject factor, cohort (first, second), and one within-subject factor, temporal structure (beat, interval), and the quantity of interest is their interaction: whether the beat−interval difference itself differs between cohorts. Because temporal structure has only two levels, that difference is a single number per participant, so the interaction can be tested directly as a two-sample comparison of those per-participant differences between cohorts, with no ANOVA to fit: under equal variances the resulting *t*, squared, is exactly the interaction *F*. We report the Welch (unequal-variance) version, which tests the same hypothesis while allowing the two cohorts to differ in spread, because the second cohort is both smaller and more variable than the first, the combination under which the pooled-variance test is anti-conservative: its participants completed uneven numbers of sessions (one or two), so their per-participant estimates carry uneven precision and inflate the cohort’s variance. Supplementary Table S2 reports four quantities, each answering a distinct question. The per-cohort effect, with its 95% confidence interval, states whether an interval−beat difference is present within a cohort and how large it is. Cohen’s *d* gives the size of the interaction itself, expressing the distance between the cohorts’ effects in pooled-standard-deviation units, on a scale comparable across tasks whose metrics differ in units. The interaction *p* tests whether the cohorts differ in that effect, but only bears against “no difference”: a non-significant value cannot distinguish a true absence of difference from a sample too small to detect one. The *Bayes Factor* BF_01_ complements it: it quantifies the evidence for “no interaction” relative to “an interaction”, which a non-significant *p* cannot.

Because only the auditory tasks changed software, the visual tasks—run in Expyriment for both cohorts— serve as a control in which the software is identical and only the sample differs. Here, the Cohort × Temporal Structure interaction was non-significant for all three tasks (Supplementary Table S2, lower panel), with small effect sizes throughout (*d* between −0.08 and +0.44). The two samples therefore behave alike where the software is held fixed, which argues against a sample difference as an explanation for the auditory comparison. Given comparable samples, the auditory comparison isolates the software change. The interval−beat effect was present in both cohorts and the Cohort × Temporal Structure interaction was non-significant for every task (Supplementary Table S2, upper panel): the robust effect of the Mean SA in the production task was reproduced in the second cohort, as were the near-null DL of the perception task and the small RT effects of the NTFD task. The interaction effect sizes were small throughout (*|d| ≤* 0.39), and smallest for the production task (*|d|* = 0.02). The Bayes factors favoured no interaction (BF_01_ between 2.1 and 3.4). At these sample sizes, even a perfectly null result would give BF_01_ *≈* 3, so the modest values reflect the second cohort’s size, not a weak signal in the data.

**Table S2.** Comparison of behavioural performance between cohorts.

| **Task** | **First cohort (*N* = 31)** | **Second cohort (*N* = 17)** | **Cohen’s *d*** | **Welch’s *t*** | ***p*** | **BF<sub>01</sub>** |
|---|---|---|---|---|---|---|
|  | **Interval−Beat [95% CI]** | **Interval−Beat [95% CI]** |  |  |  |  |
| *Auditory* |  |  |  |  |  |  |
| Production | 0.027 [0.017, 0.037] | 0.028 [0.012, 0.044] | −0.02 | −0.08 | 0.94 | 3.4 |
| Perception | −0.001 [−0.008, 0.007] | 0.010 [−0.008, 0.028] | −0.39 | −1.11 | 0.28 | 2.1 |
| NTFD | 2.9 [−1.7, 7.5] | 4.1 [−7.7, 15.8] | −0.07 | −0.20 | 0.85 | 3.3 |
| *Visual* |  |  |  |  |  |  |
| Production | 0.041 [0.025, 0.056] | 0.020 [−0.009, 0.049] | +0.44 | 1.34 | 0.19 | 1.6 |
| Perception | −0.002 [−0.016, 0.012] | 0.002 [−0.032, 0.035] | −0.08 | −0.23 | 0.82 | 3.3 |
| NTFD | 2.6 [−2.7, 8.0] | 3.7 [−4.0, 11.3] | −0.07 | −0.24 | 0.82 | 3.3 |

The condition effect (interval − beat) is given per cohort with its 95% CI. Cohen’s *d* is the standardized effect size of the Cohort × Temporal Structure interaction (the between-cohort difference in interval − beat, in pooled-standard-deviation units), tested by the Welch two-sample statistic – equivalently, whether the cohorts’ condition effects differ – and accompanied by a default-prior JZS Bayes factor BF<sub>01</sub> (evidence for no interaction; 1–3 weak, 3–10 moderate). Welch–Satterthwaite degrees of freedom ranged from 13 to 17. The auditory tasks test the re-implementation, in which the second cohort used PsychoPy; the visual tasks, run in Expyriment for both cohorts, are a same-software control isolating the sample. Metrics: *Mean Signed Asynchrony* (production, proportion of *S*), *Difference Limen* (perception, proportion), Reaction Time (NTFD, ms). The first cohort is the imaging sample.

In summary, the visual control indicates that the two cohorts are comparable samples. In the auditory comparison, reducing the timing jitter did not detectably change the behavioural differences between beat and interval sequences. Behavioural performance was therefore consistent across the two cohorts, although, given the size of the second cohort, small effects of the jitter cannot be excluded.

## Notes

### Competing Interest Statement

The authors have declared no competing interest.

